# Structure of E3 ligase E6AP with a novel proteasome-binding site provided by substrate receptor hRpn10

**DOI:** 10.1101/797027

**Authors:** Gwen R. Buel, Xiang Chen, Raj Chari, Maura J. O’Neill, Danielle L. Ebelle, Conor Jenkins, Vinidhra Sridharan, Sergey G. Tarasov, Nadya I. Tarasova, Thorkell Andresson, Kylie J. Walters

## Abstract

Regulated proteolysis by the proteasome involves ∼800 enzymes for substrate modification with ubiquitin, of which ∼600 are E3 ligases. We report here that E6AP/UBE3A is distinguished from other ubiquitin E3 ligases by having a 12 nM binding site at the proteasome contributed by substrate receptor hRpn10/PSMD4/S5a. Intrinsically disordered by itself, and previously uncharacterized, this domain in hRpn10 locks into a novel well-defined helical structure to form an intermolecular 4-helix bundle with the E6AP AZUL domain, which is unique to this E3. We thus name the h<u>R</u>pn10 <u>AZUL</u>-binding domain RAZUL. We further find in human cells that loss of RAZUL by CRISPR-based gene editing leads to loss of E6AP at the proteasome, where associated ubiquitin is correspondingly reduced, suggesting that E6AP ubiquitinates substrates at or for the proteasome. Altogether, our findings indicate E6AP to be a privileged E3 for the proteasome, with a dedicated, high affinity binding site contributed by hRpn10.

## Introduction

The 26S proteasome is a 2.5 MDa complex responsible for regulated protein degradation^1, 2^, with substrates typically ubiquitinated by a hierarchical enzymatic cascade^3^. An E1 activating enzyme charges ubiquitin to become a protein modifier and transfers it to an E2 conjugating enzyme which, in concert with an E3 ligating enzyme, subsequently attaches ubiquitin to a substrate. ∼600 E3s exist in humans that can either accept the charged ubiquitin for direct transfer to a substrate or serve as a scaffold for ubiquitin transfer from the E2 to a substrate^4–6^. Following ubiquitination, receptor sites in the proteasome contributed by Rpn1, Rpn10, and Rpn13 recognize ubiquitin directly or the ubiquitin fold of shuttle factor ubiquitin-like (UBL) domains^7–14^; shuttle factors bind ubiquitinated substrates by one or more ubiquitin-associated (UBA) domain^15–17^. At the proteasome, ubiquitin chains are hydrolyzed by deubiquitinating enzymes (DUBs) Rpn11^18^, UCHL5/Uch37^19^, and Usp14^20–22^, as the marked substrate is translocated through an ATPase ring for entry into the hollow interior of the proteolytic core particle (CP)^2, 23–25^. The integrity of the ubiquitin-proteasome pathway is essential for cellular homeostasis with dysfunction linked to disease, including cancer and neurodegeneration. Inhibitors of the CP are used to treat hematologic cancers^26–28^ and additional proteasome subunits are being pursued as synergistic targets, including hRpn13^29–34^.

Hijacking of ubiquitin E3 ligase E6AP/UBE3A by high risk human papilloma virus E6 oncoprotein contributes to cervical cancer by inducing ubiquitination and in turn degradation of tumor suppressor p53^35–37^. Moreover, loss-of-function mutations in E6AP associate with Angelman syndrome^38–40^ and elevated gene dosage with autism spectrum disorders^41^. How aberrant E6AP mechanistically contributes to these neurological diseases is an active area of investigation. It distributes in an isoform-dependent manner between the nucleus and cytosol of neurons^42, 43^ and contains an N-terminal Zn-binding AZUL (<u>a</u>mino-terminal <u>z</u>inc-binding domain of <u>u</u>biquitin E3a ligase) domain^44^ that binds to Rpn10 in the proteasome^45^, and is required for E6AP nuclear localization^43^. An Angelman syndrome-associated missense mutation in the E6AP HECT domain interferes with E6AP nuclear localization^43^, although the connection between this mutation and the requirement for the AZUL domain is not known. E6AP also stimulates Wnt/β-catenin signaling, a function that requires its ubiquitin ligase activity and interaction with the proteasome, and is disrupted by an autism-linked E6AP mutation^45, 46^.

Although multiple E3 ligases have been reported to associate with the proteasome^22, 47, 48^, no E3 ligase-proteasome complex structure is available, nor has any domain been identified for recruiting an E3 ligase to the proteasome. Here, we identify a novel region at the C-terminal end of hRpn10 that forms a binding site in the proteasome for E6AP. By using biophysical techniques including NMR spectroscopy, we find this region to be disordered when unbound, but upon binding to E6AP, to fold into an independent structural domain characterized by two helices that pack against the E6AP AZUL to form a 4-helix bundle. To test the significance of the E6AP-binding domain, we used gene editing to generate cell lines in which it is deleted and find that this hRpn10 domain contributes E6AP to the proteasome. We unexpectedly find that hRpn10 levels are coupled to E6AP cellular protein levels. Altogether, our data suggest a dual regulatory role for hRpn10 in E6AP function and that through hRpn10, E6AP is a privileged ubiquitin E3 ligase for the proteasome.

## Results

### Human Rpn10 contains a C-terminal domain that binds E6AP

Rpn10 has an N-terminal von Willebrand factor type A (VWA) domain that assembles into the proteasome and a UIM region for binding ubiquitinated proteins; these domains complete the protein in fungi (Supplementary Fig. 1a). In higher eukaryotes, Rpn10 contains an additional conserved ∼70 amino acids at the C-terminus (Fig. 1a and Supplementary Fig. 1a). We recorded a 2D NMR experiment on ^15^N-hRpn10 spanning 196-377, which encompasses the UIM region and uncharacterized C-terminal end. The resulting spectrum indicated that the UIM region, readily identified by our previous assignment of these amino acids^49^, is unperturbed by the additional C-terminal sequence (Supplementary Fig. 1b). Ubiquitin addition to ^15^N-hRpn10^196–377^ demonstrated expected shifting of the UIMs^50, 51^ but no effect for the unassigned signals (Supplementary Fig. 1c). Thus, the additional sequence does not interact with the UIM region, nor does it bind ubiquitin.

**Figure 1.**
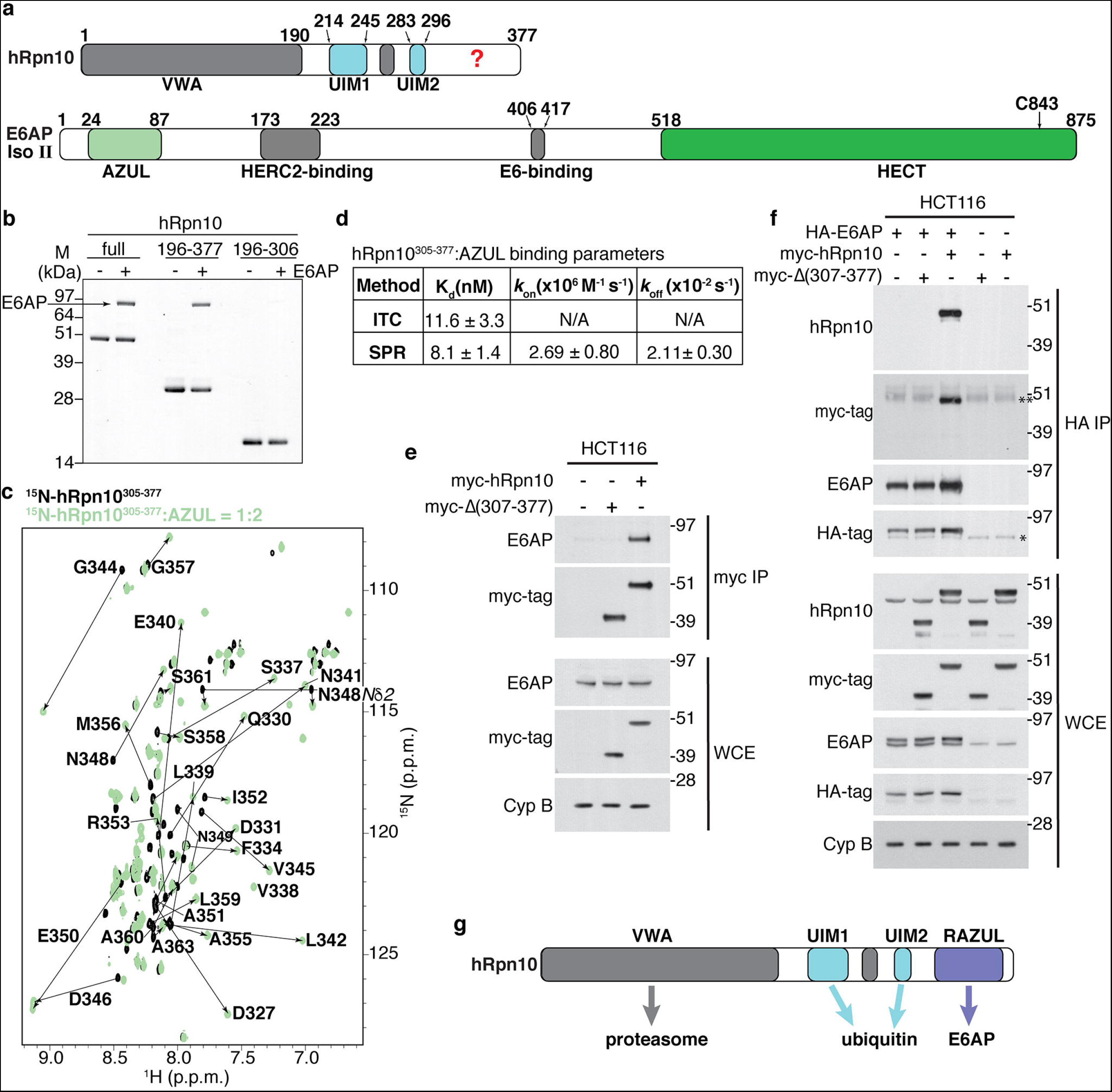
A C-terminal domain in hRpn10 binds E6AP AZUL. (**a**) Positions of known functional domains within hRpn10 (top) and E6AP isoform II (bottom). ‘?’ indicates hRpn10 uncharacterized region and the E6AP catalytic cysteine C843 is indicated. (**b**) Pull-down assay of His-tagged hRpn10^full-length^ (full), hRpn10^196–377^ or hRpn10^196–306^ without (-) or with (+) incubation of E6AP. (**c**) ^1^H, ^15^N HSQC spectra of 0.2 mM ^15^N-hRpn10^305–377^ (black) and with 2-fold molar excess unlabeled AZUL (green). Shifted signals are labeled. (**d**) Table summarizing K_d,_ *k*_on_, and *k*_off_ average values with standard deviations for the hRpn10^305–377^: AZUL interaction measured by ITC and/or SPR. N/A, not applicable. (**e**) HCT116 lysates expressing empty vector, myc-hRpn10 full length, or myc-Rpn10 with the RAZUL domain deleted (⊗307-377) were subjected to myc-immunoprecipitation with anti-myc-tag nanobody-coupled agarose. Whole cell extracts (WCE) and myc-immunoprecipitates were immunoprobed with the indicated antibodies. Cyclophilin B (Cyp B) is used as a loading control in **e**-**f**. (**f**) Lysates from HCT116 cells expressing HA-E6AP and the myc-hRpn10 constructs of **e** were subjected to HA IP followed by immunoblotting, as indicated. *, non-specific interaction; **, heavy chain antibody. (**g**) Schematic representation highlighting interaction domains of hRpn10 including newly identified RAZUL.

To identify proteins that interact directly or in complex with the C-terminal hRpn10 domain, we sub-cloned the region spanning 305-377 in frame with GST, expressed and purified the fusion protein from *E. coli*, and bound purified GST-hRpn10^305–377^ to glutathione sepharose resin for incubation with lysates from 293T (human embryonic kidney epithelial) or HCT116 (colorectal carcinoma) cells. After washing, resin-bound proteins were separated by SDS-PAGE, eluted from the gel, digested with trypsin, and analysed by mass spectrometry; parallel experiments were done with GST protein as a control. The only hit identified for either 293T or HCT116 lysate was E6AP (Supplementary Fig. 2a).

To test whether E6AP binds to the hRpn10 C-terminal region directly, we incubated full-length E6AP with Ni-NTA resin pre-bound to His-hRpn10^full-length^, His-hRpn10^196–377^, or His-hRpn10^196–306^. hRpn10^full-length^ and hRpn10^196–377^ bound E6AP, whereas hRpn10^196–306^ did not (Fig. 1b). We next added unlabelled AZUL to ^15^N-hRpn10^305–377^ and monitored the effect by 2D NMR to find hRpn10 signals shifted (Fig. 1c), indicating binding. We assigned both the free and AZUL-bound state, as described in Methods, and quantified the changes to find D327–M356 perturbed (Supplementary Fig. 2b). Analogous experiments with unlabelled hRpn10^305–377^ and ^15^N-E6AP^AZUL^, aided by previous assignments^44^, indicated residues in both AZUL helices to be significantly shifted by hRpn10 addition (Supplementary Fig. 2c and 2d). Altogether, these experiments indicate direct binding between E6AP^AZUL^ and hRpn10^305–377^, consistent with recent publications implicating AZUL and hRpn10 to E6AP interaction with the proteasome^43, 45^.

To assess the strength of hRpn10^305–377^:AZUL interaction, we used isothermal titration calorimetry (ITC) with the AZUL added incrementally to hRpn10^305–377^; a K_d_ value of 11.6 ± 3.3 nM was measured (Fig. 1d and Supplementary Fig. 2e), indicating similar strength to hRpn13 interaction with the proteasome^52, 53^. Surface plasmon resonance (SPR) similarly revealed a K_d_ value of 8.1 ± 1.4 nM for GST-hRpn10^305–377^ binding to AZUL (Fig. 1d and Supplementary Fig. 2f).

To test whether the hRpn10 C-terminal region is required for interaction with endogenous E6AP in cells, lysates from HCT116 cells expressing either myc-hRpn10^full-length^ or myc-hRpn10^1–306^ were subjected to immunoprecipitation with anti-myc nanobody-coupled agarose. Co-immunoprecipitation of E6AP was observed with full-length but not truncated hRpn10 (Fig. 1e). We further co-expressed HA-E6AP with either hRpn10 construct and immunoprecipitated E6AP with anti-HA antibodies to find co-immunoprecipitation of full-length, but not truncated, hRpn10 (Fig. 1f).

Altogether, these data indicate that the E6AP AZUL is a strong interaction partner of hRpn10^305–377^ and we henceforth refer to this domain in Rpn10 as RAZUL (<u>R</u>pn10 <u>AZUL</u> binding domain, Fig. 1g).

### E6AP protein levels depend on hRpn10

To test the significance of the RAZUL:AZUL interaction, we considered generating a full hRpn10 knockout cell line. However, we found that hRpn10 knockdown by siRNA resulted in loss of proteasome components hRpn8 and hRpn11 associating with proteasome ATPase Rpt3 (Fig. 1a), consistent with an early report that Rpn10 is necessary for base-lid interactions in the yeast proteasome^54^. To avoid such proteasome defects, we utilized CRISPR/Cas9 to generate a cell line in which hRpn10 lacks RAZUL but retains intact VWA and UIMs (see Methods). Deletion of RAZUL was assayed by immunoblotting, as demonstrated for clones 13 and 14 (Fig. 2b); we henceforth refer to these cell lines as *ΔRAZUL*.

**Figure 2.**
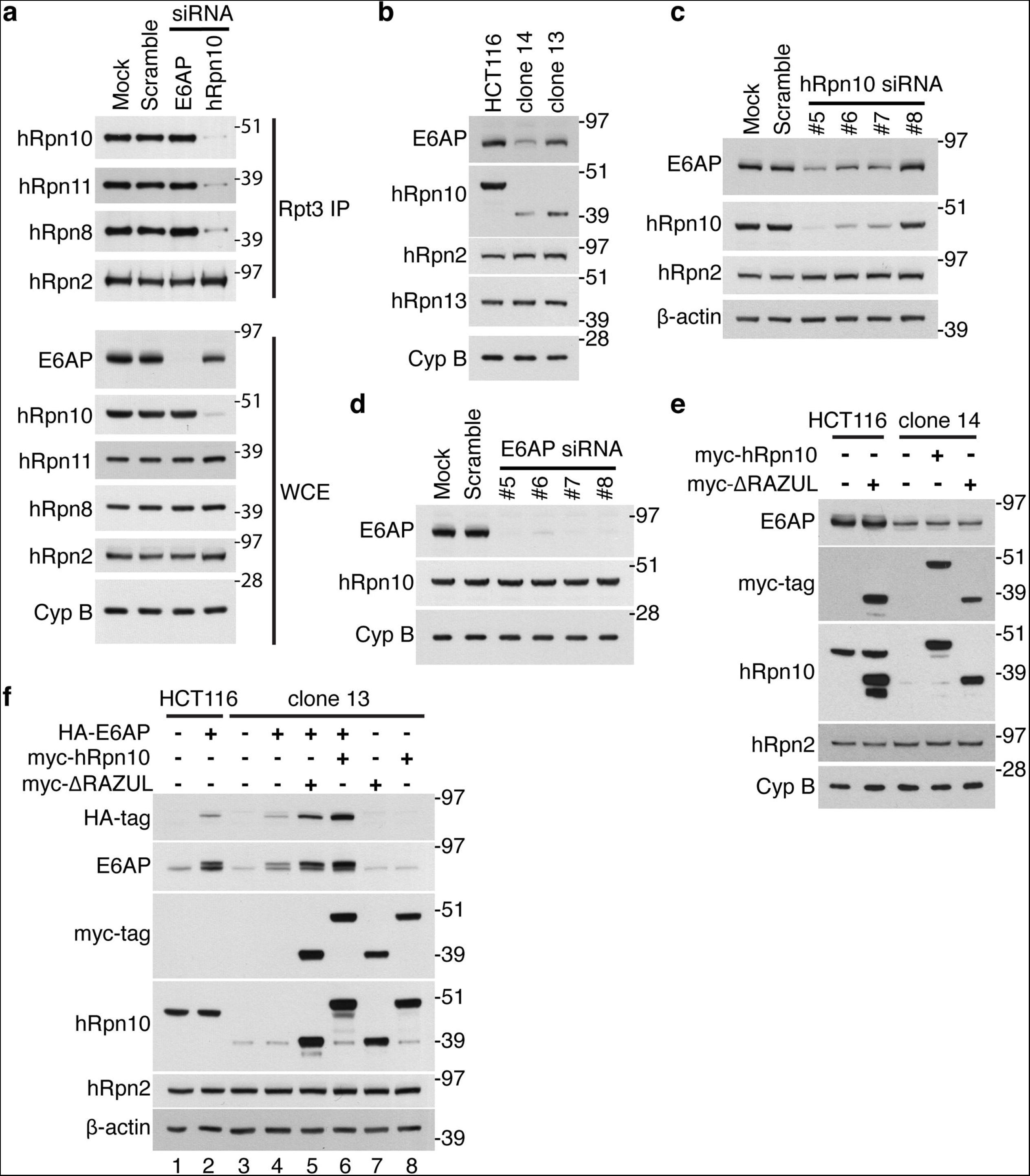
E6AP levels depend on hRpn10. (**a**) Lysates from HCT116 cells transfected with siRNAs to E6AP or hRpn10, as well as mock and scrambled control samples, were immunoprecipitated with Rpt3 antibodies. WCE and immunoprecipitates were immunoprobed as indicated. Cyclophilin B (Cyp B) is used as a loading control in **a**-**b** and **d**-**e**. (**b**) Immunoblots of parental HCT116 cells and two clonal cell lines (clone 14 and clone 13) generated by CRISPR-mediated truncation of hRpn10. (**c-d**) hRpn10 (**c**) or E6AP (**d**) was knocked down in HCT116 cells by four different siRNAs and the cell lysates immunoprobed as indicated. Mock and scrambled control samples are included. β-actin is used as a loading control in **c** and **f**. (**e**) Lysates from HCT116 or clone 14 cells expressing myc-hRpn10 constructs were immunoprobed as indicated. (**f**) Lysates from HCT116 or clone 13 cells expressing HA-E6AP and/or myc-hRpn10 constructs were immunoprobed as indicated.

Lower levels of truncated hRpn10 were consistently observed in *ΔRAZUL* cells compared to the full-length protein in *WT* cells, with clone 14 producing less protein than clone 13 (Fig. 2a, second panel). Unexpectedly, E6AP levels correlated with hRpn10 ΔRAZUL levels (Fig. 2a, top panel). To test whether the reduced protein levels are due to protein degradation, *WT* and *ΔRAZUL* cells were treated with 10 μM MG132 for four hours to inhibit the proteasome. As expected, MG132 treatment caused ubiquitinated proteins to accumulate in both cell lysates (Supplementary Fig. 3, third panel). No increase was observed however for either hRpn10 ΔRAZUL or E6AP, nor were higher molecular weight bands apparent (Supplementary Fig. 3). We assayed E6AP levels in cells with siRNA knockdown of hRpn10 to similarly find direct correlation (Fig. 2c). Loss of E6AP by siRNA treatment however had no effect on hRpn10 levels (Fig. 2d).

We attempted to rescue E6AP levels in *ΔRAZUL* cells (clone 14) by expressing hRpn10^full-length^ or ΔRAZUL protein; however, no increase in endogenous E6AP protein was observed for either condition (Fig. 2e). We further interrogated this effect in *WT* or *ΔRAZUL* cells by expressing E6AP and hRpn10 constructs either independently or in combination. Exogenous expression of E6AP was reduced in *ΔRAZUL* (clone 13) compared to *WT* cells (Fig. 2f, lane 2 versus lane 4). Exogenous E6AP levels were boosted however in ΔRAZUL cells (clone 13) when either truncated (lane 5) or full-length (lane 6) hRpn10 was co-transfected with E6AP (Fig. 2f); thus the effect is not dependent on the RAZUL:AZUL interaction, however full-length hRpn10 had a stronger effect on E6AP levels than truncated hRpn10. Altogether, these data link hRpn10 to E6AP production with possible spatial or temporal regulation that allows rescue of E6AP when the two proteins are co-expressed.

### E6AP AZUL binds to hRpn10 RAZUL at the proteasome

We used *ΔRAZUL* cells to test whether the hRpn10 RAZUL recruits E6AP to the proteasome, boosting hRpn10 levels by transient transfection. Proteasomes from lysates of *WT* and *ΔRAZUL* cells (clone 13) expressing myc-tagged hRpn10^full-length^ or ΔRAZUL protein were immunoprecipitated with Rpt3 antibodies and immunoprobed for hRpn10, E6AP, or proteasome component hRpn2 (as a control). E6AP co-immunoprecipitated with proteasomes from *WT* (Fig. 3a, lane 2 and 3), but not *ΔRAZUL* (Fig. 3a, lane 5) cells. Expression of full-length (lane 7) but not RAZUL-truncated hRpn10 (lane 6) resulted in observable E6AP co-immunoprecipitation with proteasomes isolated from *ΔRAZUL* cells (Fig. 3a); we attribute the lower amounts of E6AP co-immunoprecipitated with proteasomes of hRpn10^full-length^-expressing *ΔRAZUL* cells to the reduced abundance of endogenous E6AP in this cell line, as described above (Fig. 2a). This experiment indicates that the hRpn10 RAZUL contributes E6AP to the proteasome.

**Figure 3.**
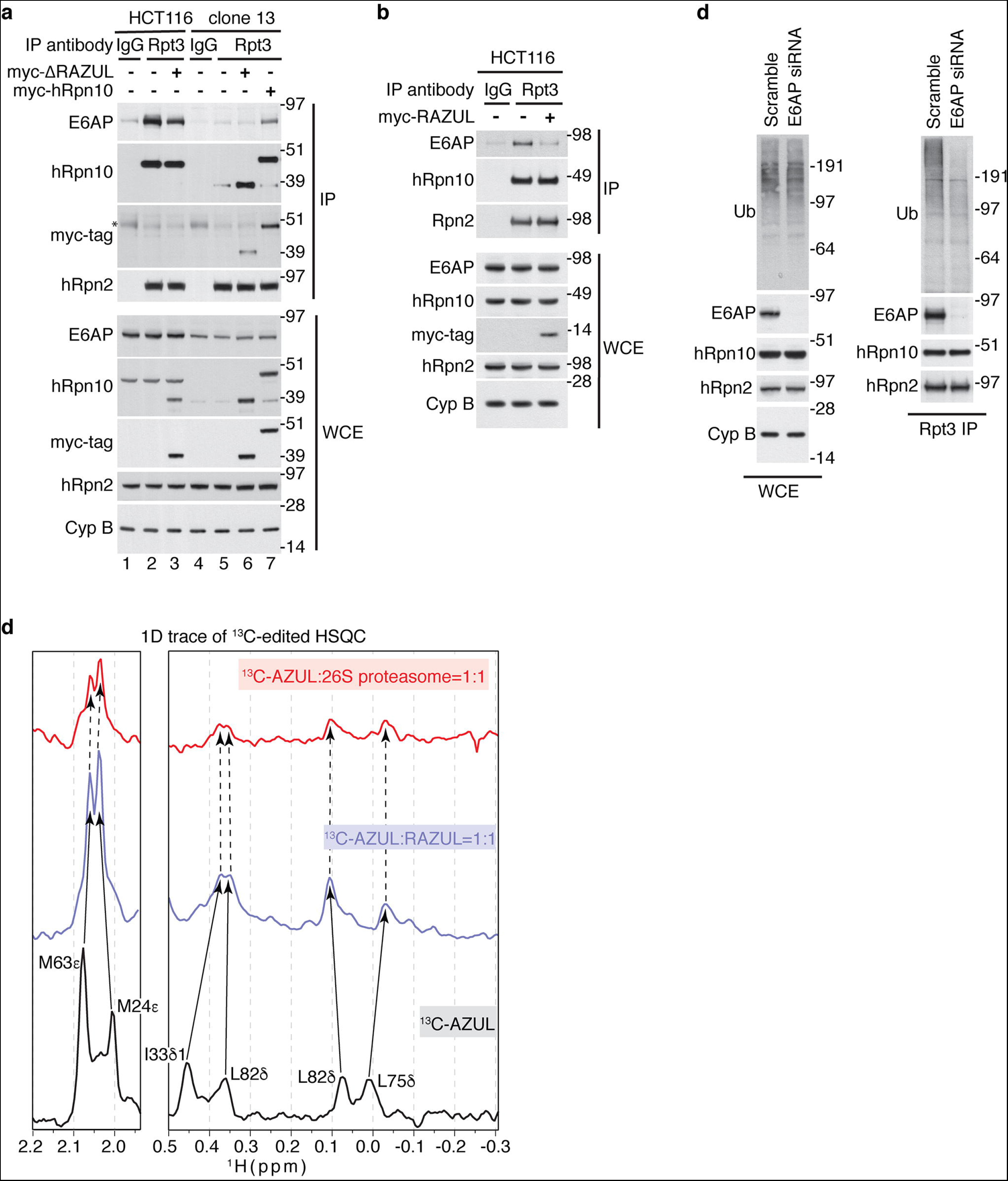
hRpn10 RAZUL contributes E6AP to the proteasome. (**a**) Immunoblots of Rpt3 immunoprecipitates or WCE from HCT116 or clone 13 lysates expressing myc-hRpn10 constructs. *, heavy chain antibody. Cyclophilin B (Cyp B) is used as a loading control for WCE samples in **a**-**c** and hRpn2 as a positive control for the immunoprecipitation. IgG controls are included. (**b**) Immunoblots of Rpt3 or IgG (control) immunoprecipitates or WCE of lysates from HCT116 cells transfected with empty vector (as a control) or myc-hRpn10 RAZUL. (**c**) Immunoblots of Rpt3 immunoprecipitates or WCE from lysates of HCT116 cells transfected with a scrambled control or siRNA against E6AP. (**d**) Selected regions from 1D ^13^C-edited, ^1^H NMR experiments acquired at 850 MHz and 25°C for free ^13^C-AZUL (black) or mixtures with equimolar unlabeled RAZUL (blue) or 26S proteasome (red). The concentration of each sample was 0.3 μM and 200,000 scans were recorded for each experiment.

We further tested whether RAZUL could compete with the proteasome for E6AP binding. Proteasomes immunoprecipitated from HCT116 cells over-expressing myc-hRpn10 RAZUL were immunoprobed for E6AP and compared to empty vector transfected cells and an IgG control. Expression of RAZUL caused E6AP to be lost from the proteasome (Fig. 3b).

To test the impact of E6AP at the proteasome, Rpt3 immunoprecipitates from lysates of cells transfected with a scrambled control or siRNA against E6AP were immunoprobed for ubiquitin. While no major change in total ubiquitin levels was observed at the level of whole cell extract (WCE), a reduction in bulk ubiquitin was apparent at the proteasome following E6AP loss, particularly for higher molecular weight species (Fig. 3c).

To test directly whether the RAZUL:AZUL interaction observed for the isolated domain complex is maintained in the intact proteasome, we added equimolar ^13^C-AZUL to RAZUL or 26S proteasome (Supplementary Fig. 4a) and acquired a 1D ^13^C-edited, ^1^H NMR experiment for comparison to free ^13^C-AZUL. Binding to RAZUL induced shifting (Supplementary Fig. 4b, middle vs bottom panel), as exemplified by methyl groups of M24 and M63 (Fig. 3d, left panel) and I33, L75, and L82 (Fig. 3d, right panel). The spectrum acquired with proteasome added closely mimicked that with only RAZUL added (Fig. 3d and Supplementary Fig. 4b), indicating that AZUL binding to RAZUL occurs identically at the proteasome.

### E6AP induces helicity in RAZUL

We used NMR to solve the structure of the RAZUL:AZUL complex, as described previously^55, 56^ and in Methods, with the data summarized in Table 1. We recorded ^13^C-half-filtered NOESY experiments on samples of the complex with one protein ^13^C labeled and the other unlabeled to measure unambiguous intermolecular interactions (Supplementary Fig. 5). Altogether, 217 interactions between AZUL and RAZUL were identified (Table 1). The 15 lowest energy structures converged with a root-mean-square deviation (r.m.s.d.) of 0.59 Å (Fig. 4a).

**Figure 4.**
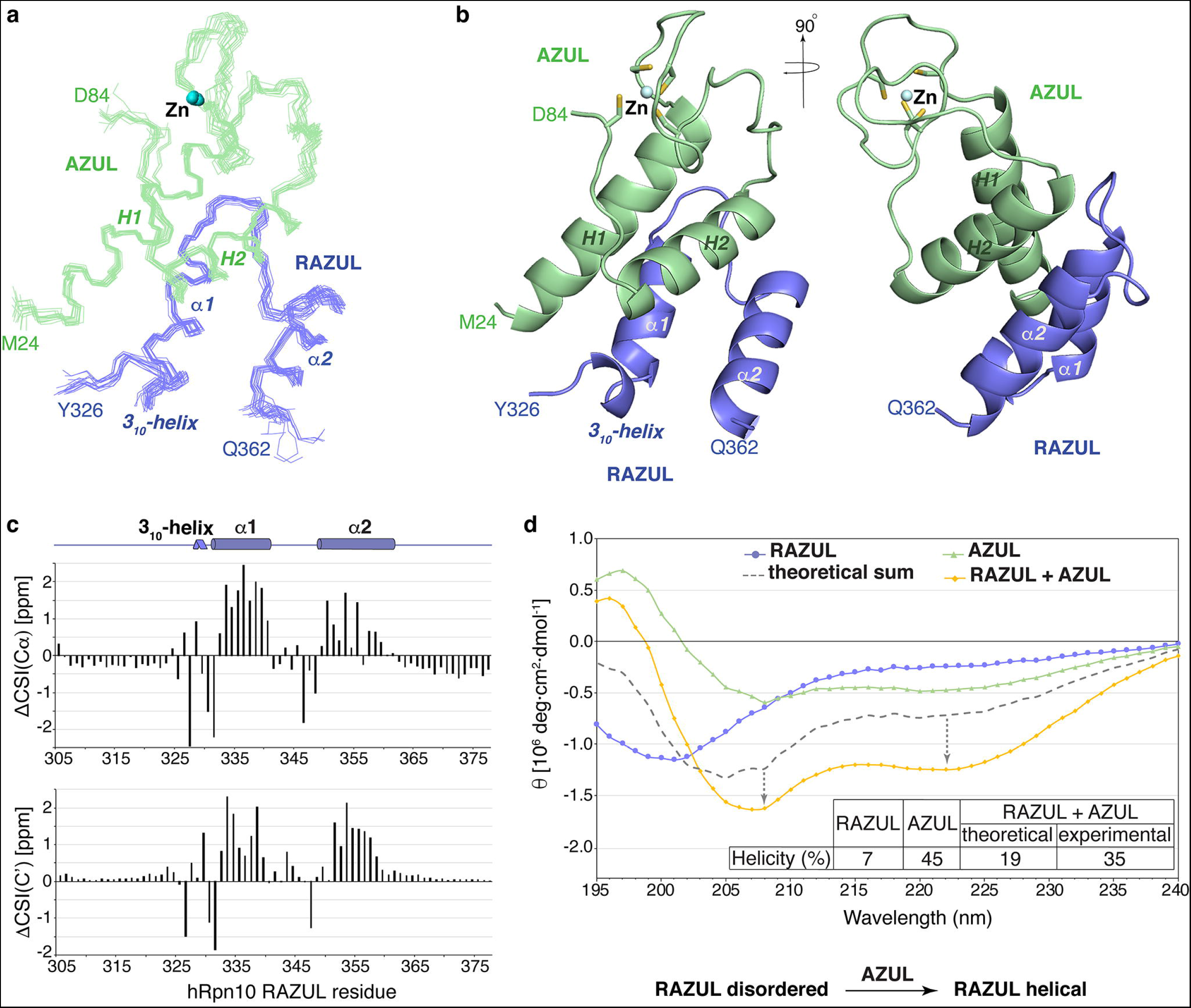
AZUL induces helicity in hRpn10 RAZUL. (**a**) Backbone trace labeling terminal residues of 15 lowest energy RAZUL (blue):AZUL (green) structures superimposing secondary structures with Zn displayed (blue sphere). Secondary structures and Zn are labeled in **a**-**b**. (**b**) Representative ribbon diagram of the RAZUL:AZUL structure with sidechain atoms of Zn-coordinating cysteines displayed as sticks with sulfur yellow. (**c**) AZUL-induced CSI change (ΔCSI) for RAZUL Cα (top) and C’ (bottom). ΔCSI(Cα)=CSI(Cα_bound_)-CSI(Cα_free_); ΔCSI(C’) = CSI(C’_bound_)-CSI(C’_free_). (**d**) CD spectra of RAZUL (blue), AZUL (green), and the mixture (orange). The theoretical sum spectrum based on free values (grey) is displayed, with a table listing predicted helicity and schematic depicting the effect of AZUL on RAZUL structure.

**Table 1.**
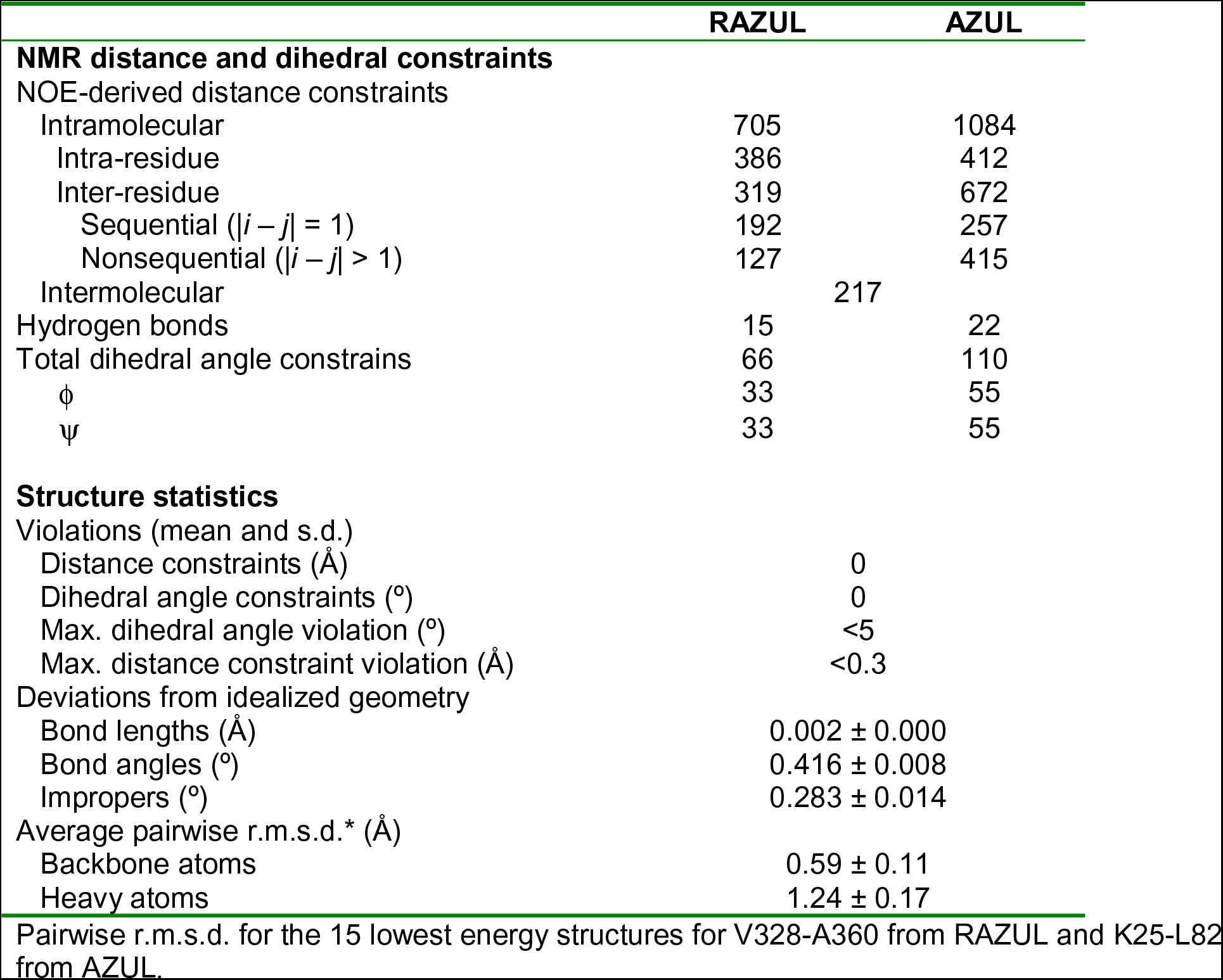
Structural statistics for the RAZUL:AZUL complex.

Unbound AZUL is comprised of two helices (H1 and H2) and a Zn finger^44^ and this architecture is maintained in the RAZUL-bound state, with a backbone r.m.s.d. between the free and complexed structures of 1.3 Å (Supplementary Fig. 6a). RAZUL binds the AZUL helices from the opposite direction compared to the Zn finger (Fig. 4b). In this complex, two α-helices are formed in RAZUL that span P332–N341 (α1) and E350–S361 (α2), with an angle of 158.5° between the two helical axes. Directly N-terminal to RAZUL α1 is a single turn of a 3_10_-helix that spans V328-Q330 (Fig. 4b). We submitted the atomic coordinates of the RAZUL:AZUL complex to the Dali server^57^ to find no similarities with either the AZUL or RAZUL domain, indicating that RAZUL is unique and not previously described.

Comparisons of our NMR data acquired on free and AZUL-bound RAZUL indicate that RAZUL acquires helicity upon binding to AZUL. Carbonyl and Cα values when compared to random coil taking into account amino acid type yields a chemical shift index (CSI) that informs on secondary structure; these values shift to reflect greater helicity for RAZUL when bound to AZUL (Fig. 4c and Supplementary Fig. 6b). Moreover, intramolecular interactions characteristic of helicity were observed following AZUL addition, but not for free RAZUL (Supplementary Fig. 6c and 6d). Overall, our NMR data indicate that RAZUL switches from a poorly ordered state to a well-defined helical state following AZUL binding. We interrogated this finding further by circular dichroism (CD) spectroscopy, as done previously for Rap80^58, 59^. CD measurements indicated 7 and 46% helicity respectively for unbound RAZUL (blue) and AZUL (green), and a theoretical spectrum (gray dashed line) for the mixture with markedly less spectral features of helicity compared to the recorded experimental spectrum (orange, Fig. 4d). 35% overall helicity is indicated from the experimental CD data recorded on the complex, consistent with the 36% helicity determined by NMR (Fig. 4b). The AZUL secondary structure is unaltered by binding to RAZUL (Supplementary Fig. 6a), leading us to conclude that the observed difference between the theoretical and experimental CD spectra reflects increased helicity for RAZUL, consistent with the NMR data (for example, Fig. 4c).

### Structure of the RAZUL:AZUL complex

At the molecular interface, a 4-helix bundle is formed by two pairs of helices from AZUL and RAZUL stacking against each other (Fig. 4b). RAZUL α1 is centered between the two AZUL helices by hydrophobic interactions involving F334, L335, V338 and L339 as well as L342 and V345 from the RAZUL α1/ α2 loop (Fig. 5a and 5b). These residues interact with A29, I33, and Y37 from AZUL H1 and A69 and L73 from AZUL H2 (Fig. 5a and 5b). From the 3_10_-helix, V328 and M329 form hydrophobic interactions with AZUL A29, L73 and Y76 (Fig. 5c), capping the hydrophobic contact surface formed by RAZUL α1. RAZUL α2 is more peripheral compared to α1, with A351, I352, A355, M356, and L359 interacting with A67, L70 and L73 from AZUL H2 (Fig. 5d).

**Figure 5.**
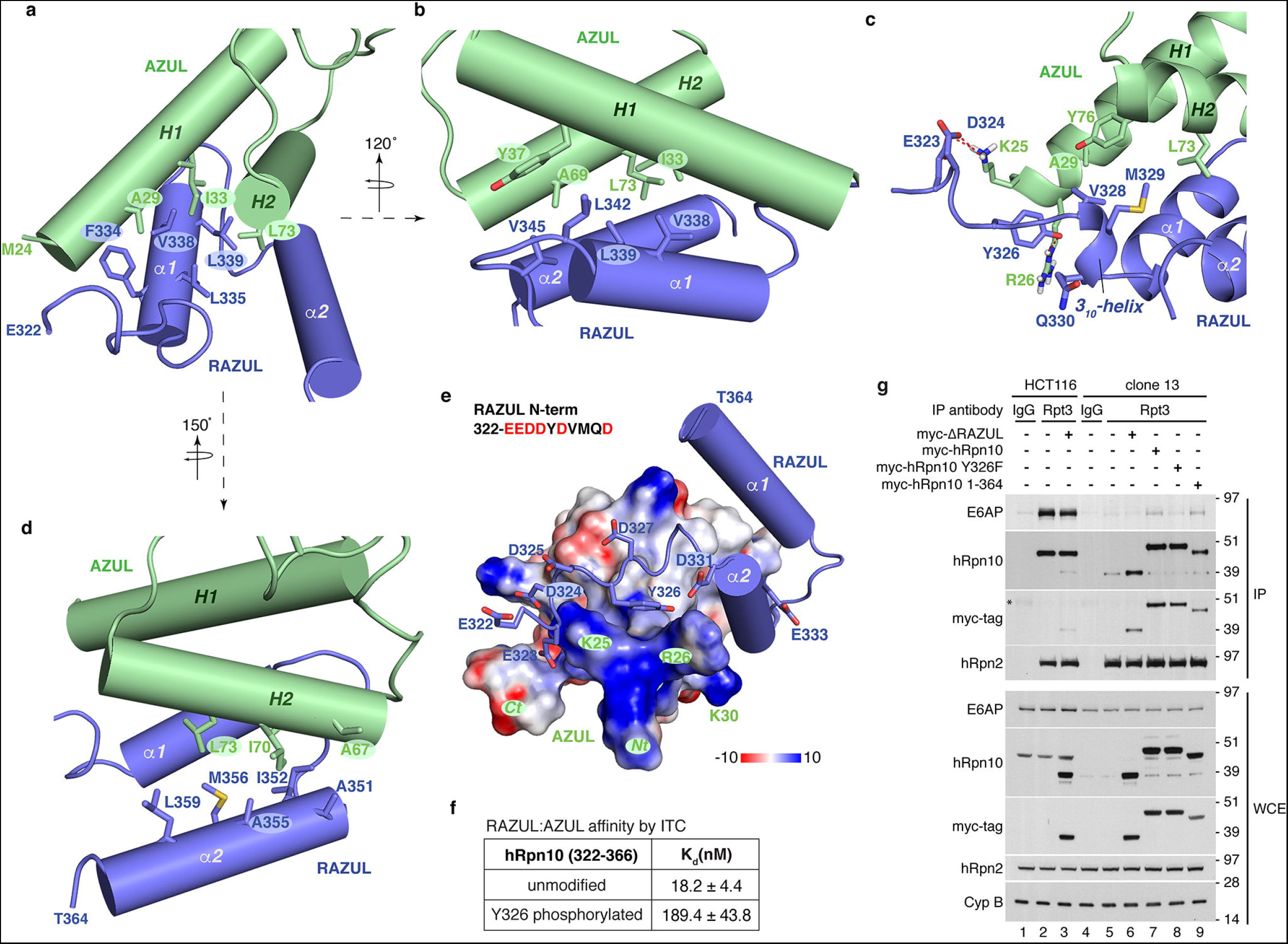
RAZUL:AZUL forms an intermolecular 4-helix bundle. (**a-d**) Regions displaying AZUL contacts with RAZUL α1 (**a**, **b**), the 3_10_ helix (**c**) and α2 (**d**) showing selected interacting sidechain atoms with oxygen, nitrogen and sulphur colored red, blue and yellow respectively. The structure in **c** is displayed as a ribbon diagram to better view the 3_10_-helix with hydrogen bonds included as red dashed lines. The structures in **a**, **b** and **d** depict helices as cylinders for simplicity. (**e**) Electrostatic surface diagram for AZUL to highlight acidic (red) and basic (blue) regions. The bound RAZUL is displayed with cylindrical helices and sticks for N-terminal acidic residues with oxygen colored red. (**f**) ITC analyses for AZUL binding hRpn10^322–366^ without or with Y326 phosphorylated. (**g**) Immunoprobing of Rpt3 immunoprecipitates or WCE from HCT116 or clone 13 lysates expressing indicated myc-hRpn10 constructs, including with RAZUL (ΔRAZUL) or the C-terminal 13 residues deleted (1-364) or with hRpn10 Y326 substituted with phenylalanine (Y326F). Cyclophilin B (Cyp B) is used as a loading control for WCE samples. *, heavy chain antibody.

The RAZUL N-terminal end (E322–D327) is rich in acidic residues (Supplementary Fig. 1a) and proximal to the positively charged AZUL N-terminal end, which includes K25 and R26 (Fig. 5e). These AZUL residues contribute three hydrogen bonds to the complex, engaging RAZUL E323, D324, and Y326 (Fig. 5c). Y326, which is phosphorylated in Jurkat cells^60^, forms a hydrogen bond with the AZUL R26 sidechain in 80% of calculated structures, as well as hydrophobic contacts with AZUL K25 and R26 (Fig. 5c). We tested whether adding a bulky phosphate group at this location could be deleterious for AZUL binding by synthesizing RAZUL peptides that span E322–D366 without and with Y326 phosphorylated and measuring affinity by ITC. This shorter wild-type peptide bound with an affinity within error of hRpn10^305–377^ (Fig. 5f and Supplementary Fig. 7a), as expected from the structure (Fig. 4a and 4b). Y326 phosphorylation of hRpn10^322–366^ reduced affinity for AZUL 10-fold (Fig. 5f and Supplementary Fig. 7b). We next tested the importance of the hydrogen bond contributed by Y326 by replacing this amino acid with phenylalanine. E6AP co-immunoprecipitation with the proteasome is reduced in *ΔRAZUL* cells expressing hRpn10 Y326F compared to those expressing the wild-type protein (Fig. 5g, lane 8 versus 7). We also found that losing the unstructured C-terminal region (365-377) from hRpn10 had no effect on E6AP co-immunoprecipitation with proteasomes by expressing hRpn10 (1-364) in *ΔRAZUL* cells (Fig. 5g, lane 9). This finding is consistent with the ITC data indicating equivalent AZUL affinity for hRpn10^322–366^ (Fig. 5f).

Altogether, these data indicate that the hRpn10 RAZUL forms an independent structural domain that contributes E6AP to the proteasome.

## Discussion

∼600 ubiquitin E3 ligases exist in humans. We report here that proteasome substrate receptor hRpn10 evolved a 12 nM affinity binding domain for recruiting E6AP to the proteasome through the N-terminal AZUL domain, which is a unique feature of E6AP. This finding provides new foundational knowledge that impacts future studies aimed at addressing the role of E6AP in cervical cancer, Angelman syndrome, and autism. Our results also redefine the current models of the 26S proteasome to include a dedicated binding domain in hRpn10 that has until now been uncharacterized despite the discovery of Rpn10 as a proteasome substrate receptor over 2.5 decades ago^61^. This domain may have remained elusive in earlier studies in part due to its disordered state when unbound and its absence in fungi.

The induced folding of RAZUL upon binding to E6AP is similar to the coupled folding reported in previous studies^62, 63^, such as the N-terminal transactivation (TAD) domain of p53, which exchanges between a disordered and partially helical conformation when unbound^64–66^ and forms a stable amphipathic α-helix when complexed with the E3 ligase Mdm2^67^. Increasing intrinsic helicity of the p53 TAD domain yields stronger binding to Mdm2 *in vitro* and in cells^68^, suggesting that RAZUL could potentially be engineered to increase E6AP occupancy at the proteasome in future work or alternatively for E6AP targeting.

In contrast to hRpn10, the other two proteasome receptors, Rpn1 and Rpn13, contribute binding sites for DUBs^11, 22, 69–71^. Proximity between substrate receptors and DUBs is no doubt conducive to ubiquitin recycling and the chain removal needed for substrate translocation into the proteolytic CP. Although it does not contribute a DUB-binding site, Rpn10 is proximal to the proteasomal DUB Rpn11^72, 73^. Why would the proteasome be benefitted by physically linking a substrate receptor to a ubiquitin E3 ligase? One possibility would be to regulate proteasome subunits, considering that E6AP is reported to ubiquitinate proteasome components, including Rpn10^74, 75^, albeit at low levels^74–76^. Rpn10 is also ubiquitinated in yeast, which is proposed to regulate its presence at the proteasome^77, 78^ and affinity for ubiquitinated proteins^79^; however, yeast lack both RAZUL and E6AP (Supplementary Fig. 1a) suggesting Rpn10 ubiquitination has redundant mechanisms or is independent of E6AP. In agreement with this, we found no evidence of altered levels or a molecular weight increase indicative of ubiquitination for hRpn10 following E6AP knockdown in HCT116 cells (Fig. 2d), although it remains possible that such activity requires induction by a specific cellular event or is cell type specific.

Another possibility is that E6AP localization to the proteasome via hRpn10 RAZUL serves in the broader cellular context to allow ubiquitin chains to be remodelled for more efficient degradation, as has been previously proposed for UBE3C/HUL5^80^. UBE3C was the first reported ubiquitin ligase to physically interact with the proteasome, with its binding site likely somewhere in the proteasome base subcomplex^80, 81^; however, its specific binding site has yet to be elucidated, and its recruitment to the proteasome appears to be assisted by structurally impaired substrate^76^.

We propose that reconfiguration of ubiquitin chains by E6AP enables more efficient degradation of proteasome substrates. Various models have been proposed to explain how different chain lengths/linkages affect the degradation rate of proteins, with multiple short ubiquitin chains shown to have higher efficiency of degradation than a single long chain^82^. Multiple ubiquitin chains on a substrate may more readily enable multiplexed points of contact with receptor sites and associated/nearby DUBs around the degradation channel and thereby enable more efficient substrate translocation. E6AP interacting with the proteasome through the Rpn10 RAZUL would be ideally situated to modify substrates in this manner, with proximity to substrates recruited by the Rpn10 UIMs or associated shuttle factors. This model is supported by our finding that E6AP knockdown reduces ubiquitin co-immunoprecipitating with the proteasome, despite largely unchanged ubiquitin levels in the whole cell extract (Fig. 3c). It has also been demonstrated that ubiquitin chains consisting of K11/K48 branched chains are more efficiently degraded by the proteasome than K11-linked chains^83^. E6AP is known to catalyze K48 linkages, however its specificity has not been studied extensively. It is possible that E6AP could add K48-linked ubiquitin (or other linkages, as yet to be determined) to existing K11-linked chains, in order to enhance degradation of substrates.

As human E6AP isoform 3 localizes to the nucleus in an hRpn10- and AZUL-dependent manner^43^, it may be that the E6AP’s function in the nucleus is primarily related to its association with hRpn10. By contrast, the decreased affinity for E6AP when RAZUL is phosphorylated at Y326 may imply that certain cellular contexts require less E6AP associated with the proteasome. For example, although we were unable to detect evidence of Y326 phosphorylation in HCT116 cells (data not shown), the identification of Y326 phosphorylation in Jukat cells (immortalized human T lymphocyte) may be significant given that immune cell activation is tightly regulated by ubiquitination^84^. E6AP has been reported to interact with Lck and Blk^85^, which are immune cell-specific tyrosine kinases at the plasma membrane involved in T cell receptor (TCR) or B cell antigen receptor (BCR) mediated activation, respectively. It is conceivable that upon TCR or BCR activation, an immune cell-specific kinase is recruited to hRpn10 via E6AP, and that subsequent phosphorylation of hRpn10 leads to reduction of E6AP at the proteasome, and in turn, delayed degradation of receptor components, allowing an elongated timeframe of activation. This mode of regulation could allow a fine-tuning of receptor activation by regulating E6AP activity at the proteasome in a very specific cellular location and context.

While future studies will elucidate the intricacies of E6AP function at the proteasome, our findings provide insight into the uniqueness of E6AP as an E3 ligase with a dedicated binding site at the proteasome and protein abundance that correlates with proteasome substrate receptor hRpn10. Our finding that E6AP binds a domain of hRpn10 not present in yeast in combination with our discovery that E6AP levels depend on hRpn10 suggests that these two proteins evolved together to have linked functions, underscoring the importance of E6AP at the proteasome. These results provide a new foundation towards understanding the role of E6AP in its associated disease states.

## Methods details

### Protein sample preparation for biophysics experiments

Human Rpn10^305–377^ or its amino acid substituted variants were sub-cloned into pGEX-6P-3 vector and expressed in *Escherichia coli* strain Rosetta 2 (DE3) as fusion proteins with N-terminal glutathione S-transferase (GST) and a PreScission protease cleavage site. Cells were grown at 37°C to OD_600_ of 0.5 – 0.6 and protein expression induced overnight at 17°C by addition of 0.4 mM IPTG. The cells were frozen in liquid nitrogen and stored at −80°C for ∼4 hrs, followed by resuspension in buffer 1 (50 mM Tris at pH 7.2, 300 mM NaCl, 5 mM DTT) supplemented with protease inhibitor cocktail tablets (Roche Diagnostics). Cells were lysed by sonication and spun down at 27,000 g for 30 min, after which the supernatant was incubated with pre-washed glutathione sepharose resin for 3 hrs. After extensive washing in buffer 1, hRpn10^305–377^ or its amino acid substituted variants were either separated from GST and the resin by cleaving with PreScission protease in buffer 2 (10 mM MOPS at pH 6.5, 50 mM NaCl, 5 mM DTT, 10 μM zinc sulphate), or eluted from the resin with the GST-tag intact by using buffer 3 (50 mM Tris at pH 7.2, 50 mM NaCl, 5 mM DTT, 20 mM glutathione). Further purification was achieved by size exclusion chromatography on an FPLC system ÄKTA pure (GE Healthcare) using a HiLoad 16/600 Superdex 75 (for samples with no GST-tag) or Superdex 200 (for GST-tagged protein) prep grade column in buffer 2 or 3.

E6AP^24–87^ was sub-cloned into the pET28a vector and expressed in *Escherichia coli* strain BL21 (DE3) as a fusion protein with an N-terminal polyhistidine tag and a thrombin cleavage site. Cells were grown at 37°C to OD_600_ of 0.5 – 0.6 and protein expression induced at 17°C overnight by 0.4 mM IPTG. At the time of induction, zinc sulphate was added to a final concentration of 20 μM. The cells were frozen in liquid nitrogen and stored at −80°C for ∼4 hrs, followed by resuspension in buffer 4 (10 mM MOPS at pH 7.2, 300 mM NaCl, 5 mM 21Zmercaptoethanol, 10 uM zinc sulphate) supplemented with protease inhibitor cocktail tablets (Roche Diagnostics). Cells were lysed by sonication and spun down at 27,000 g for 30 min. The supernatant was incubated with pre-washed Ni-NTA agarose resin (Qiagen) for 1 hr. After extensive washing in buffer 4, E6AP^24–87^ was separated from the His-tag and the resin by cleaving with thrombin in buffer 5 (10 mM MOPS at pH 6.5, 50 mM NaCl, 5 mM 21Zmercaptoethanol, 10 μM zinc sulphate). Further purification was achieved by size exclusion chromatography on an FPLC system ÄKTA pure (GE Healthcare) using a HiLoad 16/600 Superdex 75 prep grade column in buffer 5.

hRpn10^196–377^ was sub-cloned into the pET14b vector with an N-terminal polyhistidine tag and a thrombin cleavage site. pET11d/His_6_-hRpn10^full-length^ is a gift from Dr. Fumio Hanaoka. pET26b/His_10_-hRpn10^196–306^ was reported previously^12, 50^. N-terminal His-tagged hRpn10^full-length^, hRpn10^196–306^, and hRpn10^196–377^ were expressed from *Escherichia coli* strain BL21 (DE3) and purified in an identical manner as N-terminal His-tagged E6AP^24–87^, but eluted from the resin with the His-tag intact by using elution buffer 6 (10 mM MOPS at pH 7.2, 50 mM NaCl, 5 mM 21Zmercaptoethanol, and 250 mM imidazole), and further purified by size exclusion chromatography on an FPLC system ÄKTA pure (GE Healthcare) with a HiLoad 16/600 Superdex 75 prep grade column in buffer 5.

^15^N ammonium chloride, ^13^C glucose, and ^2^H_2_O were used for isotope labeling. All NMR samples were validated by mass spectrometry. 26S proteasome (human) was purchased (Enzo Life Sciences, Inc. BML-PW9310).

### Peptide synthesis

hRpn10^322–366^ peptide without or with Y326 phosphorylated was synthesized on a Liberty Blue Microwave peptide synthesizer (CEM Corporation) using Fmoc chemistry. To avoid oxidation, Met residues in the sequence of HVR were substituted by isosteric norleucine. The following modifications were introduced to the published protocol for high efficiency peptide synthesis^86^. The coupling with N,N’-diisopropylcarbodiimide (DIC)/ethyl 2-cyano-2-(hydroxyimino)acetate (OXYMA) was performed for 4 min at 90°C for all residues except Cys and His, for which the reaction was carried out for 10 min at 50^°^C. Removal of the Fmoc group was conducted at 90°C for 2 min for sequences containing no Cys or Asp. All deprotection cycles after Asp and Cys were conducted at room temperature to avoid racemization and aspartimide formation. Low loading Rink Amide MBHA resin (Merck) was used for the synthesis of amidated peptides and Wang resins were used for the synthesis of peptides with free carboxy-termini. The peptides were cleaved from the resin and deprotected with a mixture of 90.0% (v/v) trifluoroacetic acid (TFA) with 2.5% water, 2.5% triisopropyl-silane, 2.5% 2,2′-(ethylenedioxy)diethanethiol and 5% thioanisol. Peptides were purified on a preparative (25 mm x 250 mm) Atlantis C3 reverse phase column (Agilent Technologies) in a 90 min gradient of 0.1% (v/v) trifluoroacetic acid in water and 0.1% trifluoroacetic acid in acetonitrile, with a 10 mL/min flow rate. The fractions containing peptides were analyzed on an Agilent 6100 LC/MS spectrometer with the use of a Zorbax 300SB-C3 PoroShell column and a gradient of 5% acetic acid in water and acetonitrile. Fractions that were more than 95% pure were combined and freeze dried.

### GST-pulldown experiment and sample preparation for mass spectrometry

HCT116 and 293T cells were purchased from American Type Culture Collection (ATCC CCL-247 and CRL-3216). 293T cells were grown in DMEM (GlutaMAX^TM^-1 with 4.5 g/L D-glucose and without sodium pyruvate, Thermo Fisher Scientific) and HCT116 cells were grown in McCoy’s 5A modified medium (ATCC 30-2007), with both media supplemented with 10% fetal bovine serum (Atlanta Biologicals, Inc. S12450). Cells were grown in a 37°C humidified atmosphere of 5% CO_2_. Cells were collected, washed twice with PBS, and lysed in 1% Triton-TBS buffer (50 mM Tris at pH 7.5, 150 mM NaCl, 1 mM PMSF), supplemented with protease inhibitor cocktail (Roche Diagnostics). Total protein concentration in the lysate was determined by a Pierce bicinchoninic acid protein assay kit (Thermo Fisher Scientific 23225). 2 nmol of purified GST-tagged hRpn10^305–377^ or GST protein (Thermo Scientific 20237) was added to 20 μL of pre-washed glutathione sepharose resin for 3 hrs, washed once with buffer 7 (50 mM Tris at pH 7.5, 150 mM NaCl, 2 mM DTT, 0.5% (v/v) Triton-X100, 10 μM zinc sulphate), and the resin next incubated with 1.2 mL of cell lysate for 3 hrs. Unbound protein was removed by washing three times in buffer 8 (20 mM Tris at pH 7.6, 150 mM NaCl, 2 mM DTT, 10% glycerol, 10 μM zinc sulphate), after which resin-bound proteins were eluted, subjected to electrophoresis on a 4-12% NuPAGE Bis-Tris gel (Thermo Fisher Scientific NP0322), and visualized by Coomasie staining.

For each of the six lanes (GST-hRpn10^305–377^ or GST protein incubated with 293T or HCT116 cell lysate), the region above 39 kDa was cut into 12 bands that were placed individually into 1.5 mL eppendorf tubes. Each gel band was then further cut into small pieces and destained using 50% acetonitrile/25 mM NH_4_HCO_3_ at pH 8. After removal of the organic solvent, gel pieces were dried by vacuum centrifugation for 1 hr. Trypsin (20 ng/µL) in 25 mM NH_4_HCO_3_ at pH 8, was added to each sample (50 µL) and incubated on ice for 1 hr. 25 mM ammonium bicarbonate was added to completely saturate the bands for overnight incubation at 37°C. Peptides were extracted in 70% acetonitrile and 5% formic acid using bath sonication and supernatant solutions were dried in a speed vacuum. Samples were desalted utilizing Pierce C18 spin columns (Thermo Fisher Scientific), dried, and resuspended in 0.1% TFA prior to mass spectrometry analysis.

### Nanoflow Reversed Phase Liquid Chromatography (nanoRPLC)-Tandem Mass Spectrometry

Peptides were analyzed on a Q Exactive Hybrid Quadrupole-Orbitrap mass spectrometer (Thermo Fisher Scientific). The desalted tryptic peptide was loaded onto an Acclaim PepMap 100 C18 LC column (Thermo Fisher Scientific) utilizing a Thermo Easy nLC 1000 LC system (Thermo Fisher Scientific) connected to the Q Exactive mass spectrometer. The peptides were eluted with a 5-48% gradient of acetonitrile with 0.1% formic acid over 55 min with a flow rate of 300 nL/min. The raw MS data was collected and analyzed in Proteome Discoverer 2.2 (Thermo Fisher Scientific) with Sequest HT software and was searched against the Human Proteome database.

### His pull-down assays

His pull-down assays were performed as described previously^11, 87^. Briefly, 2 nmol of purified His-tagged hRpn10^full-length^, hRpn10^196–377^, or hRpn10^196–306^ was added to 20 μL of pre-washed Ni-NTA agarose resin (QIAGEN) for 2 hrs and washed once with buffer 5. The resin was then incubated with 200 pmol of E6AP (UBPBio K1411) for 1 hr and unbound protein removed by extensive washing with buffer 5. Resin-bound proteins were eluted and subjected to SDS-PAGE followed by visualization with Coomasie staining.

### ITC and SPR binding affinity experiments

ITC was performed at 25°C on a MicroCal iTC200 system. hRpn10^305–377^, E6AP^24–87^, and hRpn10^322–366^ peptide without or with Y326 phosphorylated were dialyzed extensively against buffer 2. Eighteen 2.1 μL aliquots of 0.462 mM E6AP^24–87^ were injected at 1000 rpm into a calorimeter cell (volume 200.7 μL) that contained 0.0405 mM hRpn10^305–377^. For measuring interaction between hRpn10^322–366^ peptides and E6AP^24–87^, eighteen 2.1 μL aliquots of 0.110 mM E6AP^24–87^ were injected at 1000 rpm into a calorimeter cell (volume 200.7 μL) that contained 0.01 mM hRpn10^322–366^ without or with Y326 phosphorylated. Blank experiments were performed by replacing protein samples with buffer and this blank data was subtracted from the experimental data during analysis. The integrated interaction heat values were normalized as a function of the molar ratio of E6AP^24–87^ to hRpn10^305–377^ or to hRpn10^322–366^ peptides, and the data were fit with MicroCal Origin 7.0 software. Binding was assumed to be at one site to yield the binding affinity Ka (1/Kd), stoichiometry, and other thermodynamic parameters.

Surface plasmon resonance experiments were recorded for GST-tagged hRpn10^305–377^ and E6AP^24–87^ with a Biacore T200 system (GE Healthcare). Utilizing the GST capture kit, 4000 RU of anti-GST antibody was covalently immobilized on a CM5 chip via amine coupling. GST-tagged hRpn10^305–377^ was then added to FC2 to a final response of 800 RU. As a negative control, GST was added to FC1 to the same response. E6AP^24–87^ was prepared in degassed, filtered HBS-P+ (GE Healthcare) buffer with 10 μM zinc sulphate. Single cycle kinetic experiments were performed using five injections (30 µL/min) of increasing concentration of protein (5–250 nM) passed over the sensor chip for 150 sec association, followed by a 420 sec dissociation. The experiments were repeated in triplicate. Buffer and reference subtracted kinetic constants (*k*_on_ and *k*_off_) and binding affinities (K_d_) were determined utilizing the Biacore T200 evaluation software (GE Healthcare).

### CD experiments

Far-UV range CD spectra (240-190 nm) of 10 μM hRpn10^305–377^, 10 μM E6AP^24–87^, the mixture of 10 μM hRpn10^305–377^ and 10 μM E6AP^24–87^, and buffer 9 (10 mM MOPS, 10 mM NaCl, 10 mM DTT, 10 μM zinc sulphate at pH 6.5, as a control) were recorded on a Jasco J-1500 circular dichroism spectrometer (Tokio, Japan) using a quartz cuvette with 1.0 mm path length and temperature controlled at 25 ± 0.1°C. All spectra were collected continuously at a scan speed of 20 nm/min and averaged over accumulation of three spectra. The buffer spectrum was subtracted from the protein spectra during data analyses. The molar ellipticity θ (in deg cm^2^ dmol^−1^) was calculated from the measured machine units m° in millidegrees at wavelength λ using the Equation 1.

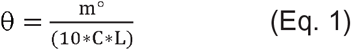

C is the concentration of the sample in mol L^-1^ and L is the path length of the cell (cm). Secondary structure analysis was conducted with the program CONTIN^88^ at DichroWeb server^89, 90^ by using the reference dataset SMP180 (190-240 nm)^91^.

### Cell culture, plasmids, siRNAs, transfections

All cell lines were grown in McCoy’s 5A modified medium (Thermo Fisher Scientific 16600082), with 10% fetal bovine serum (Atlanta Biologicals, Inc. S12450) at 37°C and 5% CO_2_. The HCT116 cell line was purchased from the ATCC (CCL-247). Myc-tagged hRpn10 constructs were generated by Genscript by inserting hRpn10 (NM_002810.2) full-length coding sequence or DNA encoding for residues 1-306 between the KpnI and XhoI restriction sites of pcDNA3.1(+)-N-Myc. The resulting constructs included an N-terminal myc tag with a Gly-Thr linker between the myc tag and hRpn10 due to the KpnI site. p4054 HA-E6AP isoform II was purchased from Addgene (plasmid #8658), originally constructed in the laboratory of Peter Howley^92^. Plasmid transfections (48 hrs) were carried out with Lipofectamine 3000 (Thermo Fisher Scientific). siRNA transfections (72 hrs) were performed with Lipofectamine RNAiMAX (Thermo Fisher Scientific). Conditions labeled ‘mock’ received only RNAiMAX, conditions labeled ‘scramble’ were transfected with ON-TARGETplus non-targetting siRNA #2 (Dharmacon D-001810-02). hRpn10 and E6AP siRNAs used were ON-TARGETplus siRNAs (Dharmacon 011365 and 005137, respectively). Where only one siRNA from the set was used, the siRNAs were 011365-05 and 005137-05.

### CRISPR-Cas9 generation of ***Δ****RAZUL*

Candidate sgRNAs flanking the starting portion of the RAZUL domain were identified using sgRNA Scorer 2.0^93^. Oligonucleotides for each sgRNA were annealed and ligated into a vector carrying Cas9-2A-mCerulean. The Cas9-2A-mCerulean was generated by digesting the pX458 backbone and replacing eGFP with mCerulean. pX458 was a gift from Feng Zhang (Addgene plasmid # 48138; http://n2t.net/addgene:48138; RRID:Addgene_48138)^94^. Plasmid donor used as template for homology-directed repair was generated using isothermal assembly of the left and right homology arms with the P2A-puromycin cassette. The left and right homology arms were amplified using PCR from 293T genomic DNA and mutations were introduced in the PAM sequence of each target site to prevent editing of the dsDNA donor. Sequences of all oligomers used are provided in Table S1. HCT116 cells were co-transfected with the Cas9-2A-mCerulean constructs in combination with the donor construct. Two days post transfection, cells were split into a fresh 6-well plate and grown in media with puromycin for two weeks. A few cell colonies were visible after two weeks that were picked using sterile tips and added to a 24-well plate with fresh media. Individual clones were then analyzed by immunoblotting using hRpn10 antibody to identify clones containing the truncation. Clones 13 and 14 were selected based on this analysis, and genomic DNA isolated from clones 13 and 14 were used in PCR amplification using primers outside the homology arms. PCR fragments of the expected size were generated, and PCR using one primer outside the homology arm and one to puromycin validated insertion of the donor sequence. PCR-amplified genomic DNA was cloned into the pCR™4-TOPO® vector using a TOPO™ TA Cloning™ Kit (Life Technologies K457501) and sequence verified.

### Cell lysis and immunoprecipitations

All cells were washed twice in cold PBS (Thermo Fisher Scientific) prior to harvesting. Cells used for Rpt3 IP were harvested on ice in 1% Triton X-100 buffer (1% Triton X-100, 50 mM Tris pH 7.5, 150 mM NaCl, 1 mM PMSF, 5 μg/mL Pepstatin A, and Roche EDTA-free protease inhibitor cocktail). Cell extracts were centrifuged for 15 min at 4°C and 20,000g, and the supernatant was isolated. IPs were performed overnight at 4°C with 1-1.5 mg total protein lysate, using 4 μL Rpt3 antibody (abcam ab140515) per condition. Protein G sepharose beads (GE healthcare 17-0618-01) were added to IPs for 3 hrs, and precipitates were washed 5-7 times with 1% Triton X-100 buffer. Immune complexes were heated to 95°C for 10 min in denaturing sample buffer prior to subjection to SDS-PAGE.

Cells used for myc-trap IPs were harvested on ice in 0.5% NP-40 lysis buffer (0.5% Nonidet P-40, 50 mM Tris pH 7.5, 150 mM NaCl, 1 mM PMSF, 5 μg/mL Pepstatin A, 10 mM sodium pyrophosphate, 10 mM NaF, 1 mM Na_3_VO_4_, and Roche EDTA-free protease inhibitor cocktail). Cell extracts were placed on ice for 30 min with extensive pipetting every 10 min. Extracts were centrifuged at 20,000g for 30 min at 4°C. Supernatants were diluted 1:1 with myc-trap dilution/wash buffer (50 mM Tris pH 7.5, 150 mM NaCl, 1 mM PMSF, 5 μg/mL Pepstatin A, 10 mM sodium pyrophosphate, 10 mM NaF, 1 mM Na_3_VO_4_, and Roche EDTA-free protease inhibitor cocktail), and 1-1.5 mg total protein lysate was incubated with 25 μL myc-trap agarose (nanobody coupled) beads (chromotek) overnight at 4°C. Nanobody-myc complexes were washed 5 times with myc-trap dilution/wash buffer and heated to 95°C for 10 min in denaturing sample buffer prior to subjection to SDS-PAGE.

In experiments not involving immunoprecipitation, cells were either harvested in 1% Triton X-100 buffer or 1% NP-40 buffer (1% Nonidet P-40, 25 mM Tris pH 7.2, 137 mM NaCl, 10% glycerol, 1 mM DTT, 5 mg/mL Pepstatin A, 1 mM PMSF, and Roche protease inhibitor cocktail). Lysates were centrifuged at 20,000g and 4°C for 15 min and supernatants were isolated for immunoblotting.

### SDS-PAGE, immunoblots, and antibodies

Protein lysates were subjected to SDS-PAGE on 4-12% NuPAGE Bis-Tris gels (Thermo Fisher Scientific NP0322) using MOPS SDS running buffer (Thermo Fisher Scientific NP0001), except in the case of Fig. 3b, where MES SDS running buffer (Thermo Fisher Scientific NP002) was used to achieve better resolution of myc-RAZUL. Proteins were transferred to 0.45 μm PVDF membranes (Thermo Fisher Scientific LC2005) using NuPAGE transfer buffer (Thermo Fisher Scientific NP00061) supplemented with 10% methanol. Following transfer, membranes were blocked with 5% milk in tris-buffered saline with 1% tween 20 (TBS-T). Blocked membranes were incubated with primary antibodies (diluted in 5% milk in TBS-T) overnight. Membranes were washed 5 times in TBS-T and incubated with HRP-conjugated secondary antibodies (diluted in 5% milk in TBS-T) for 2 hrs. Following another set of 5 washes, blots were developed using HyGlo quickspray chemiluminescent HRP detection reagent (Denville Scientific Inc. E2400) and HyBlot CL Autoradiography Film (Denville Scientific Inc. E3018). Primary antibodies used were: β-actin (Cell Signaling Technologies 4970), Cyclophilin B (Abcam ab178397), E6AP (MilliporeSigma E8655), HA-tag (MilliporeSigma H6908), myc-tag (Cell Signaling Technologies 2278), Rpn10 (Novus Biologicals NBP2-19952), Rpn2 (Bethyl Laboratories A303-851A), Rpn13 (Abcam ab140515), Rpn11 (Cell Signaling Technologies 4197), Rpn8 (abcam ab140428), and ubiquitin (MilliporeSigma MAB1510). Secondary antibodies used were: Rabbit HRP (Life technologies A16110) and mouse HRP (MilliporeSigma A9917). For blots in which a protein ran close to the heavy chain in an IP condition, Native Rabbit HRP (MilliporeSigma R3155) was used.

### NMR samples and experiments

Three NMR samples were prepared, including 1) 0.5 mM ^15^N, ^13^C, 35% ^2^H-labeled Rpn10^305–377^ mixed with unlabeled E6AP^24–87^ at 1.5-fold molar excess; 2) 0.5 mM ^15^N, ^13^C E6AP^24–87^ mixed with unlabeled Rpn10^305–377^ at 1.5-fold molar excess; 3) 0.5 mM ^15^N, ^13^C, 70% ^2^H-labeled Rpn10^196–377^. For assignment of Rpn10^305–377^ in the E6AP-bound state, or E6AP in the Rpn10^305–, 377–^bound state, 2D ^1^H-^15^N HSQC and ^1^H-^13^C HSQC, 3D HNCACB/CBCA(CO)NH, HNCO/HN(CA)CO, HCCH-TOCSY, CCH-TOCSY, ^15^N (120 ms mixing time) and ^13^C (80 ms mixing time) edited NOESY-HSQC spectra were recorded on samples 1 and 2. These experiments were also recorded on sample 3 to obtain free state assignments for Rpn10^196–377^. Unambiguous intermolecular distance constraints were obtained by using 3D ^13^C-half-filtered NOESY experiments (100 ms mixing time) recorded on samples 1 and 2. Chemical shift assignment of E6AP^24–87^ in the free state was available from a previous study^44^.

All NMR experiments were conducted at 25**°**C in buffer 10 (10 mM MOPS at pH 6.5, 50 mM NaCl, 5 mM DTT, 10 μM zinc sulphate, 1 mM pefabloc, 0.1% NaN_3_ and 5% ^2^H_2_O / 95% ^1^H_2_O), except for 2D ^1^H-^13^C HSQC, 3D HCCH-TOCSY, CCH-TOCSY, ^13^C-edited NOESY-HSQC and ^13^C-half-filtered NOESY experiments, which were acquired on samples dissolved in ^2^H_2_O. Spectra were recorded on Bruker AvanceIII 600, 700, 800, or 850 MHz spectrometers equipped with cryogenically cooled probes.

All NMR data processing was performed with NMRpipe^95^ and spectra were visualized and analyzed with XEASY^96^. Secondary structure was assessed by comparing chemical shift values of Cα and C’ atoms to random coil positions to generate a chemical shift index (CSI)^97^ and also by the TALOS+ program^98^.

### NMR titration experiments

^1^H, ^15^N HSQC experiments were recorded on 0.2 mM ^15^N-labeled samples (hRpn10^305–377^ or E6AP^24–87^) with increasing molar ratio of unlabeled ligand (E6AP^24–87^ or hRpn10^305–377^), as indicated. The amide nitrogen and hydrogen chemical shift perturbations (CSP) were mapped for each amino acid according to Equation 2.

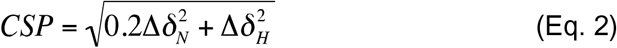

Δ*δ*_H_, change in amide proton value (in parts per million); Δ*δ*_N_, change in amide nitrogen value (in parts per million).

### 1D ^13^C-edited, ^1^H NMR experiments

Three NMR samples were prepared in buffer 11 (50 mM d^11^-Tris at pH 7.5, 50 mM NaCl, 5 mM MgCl_2_, 1.5 mM ATP-γS, 10 μM zinc sulphate, 2 mM DTT, 0.5 mM pefabloc and 5% ^2^H_2_O / 95% ^1^H_2_O), including 0.3 μM of free ^13^C labeled AZUL domain, 0.3 μM of ^13^C labeled AZUL domain mixed with equimolar unlabeled RAZUL, and 0.3 μM of ^13^C labeled AZUL domain mixed with equimolar human 26S proteasome (Enzo Life Sciences, Inc., BML-PW9310). 200,000 1D traces of a ^1^H, ^13^C HSQC experiment^99, 100^ were averaged for each sample at 25**°**C and 850 MHz with a cryogenically cooled probe.

### Structure calculation of the hRpn10 RAZUL: E6AP AZUL complex

XPLOR-NIH 2.50^101^ was used on a Linux operating system to solve the complexed structure by using NOE and hydrogen bond constraints as well as backbone ϕ and ψ torsion angle constraints derived from TALOS+^98^ (Table 1). Hydrogen bonds were generated by using secondary structure assignments and NOE connectivities with defined distances from the acceptor oxygen to the donor hydrogen and nitrogen of 1.8-2.1 Å and 2.5-2.9 Å, respectively. Hydrogen bonds restraints were not included in the initial calculation, but were in the final round of structure calculations. When calculating the structures of hRpn10 RAZUL:E6AP AZUL, intermolecular distance constraints determined from the 3D ^13^C-half-filtered NOESY experiments were used, in addition to intramolecular constraints for hRpn10 RAZUL and E6AP AZUL that were generated from ^15^N or ^13^C NOESY spectra acquired on the complexes (Table 1). The complexed structures were calculated from 50 linear starting structures of hRpn10 RAZUL and E6AP AZUL molecules, which were subjected to 2,000 steps of initial energy minimization to ensure full spatial sampling and appropriate coordinate geometry. The structures were next confined according to the inputted data by subjecting them to 55,000 simulated annealing steps of 0.005 ps at 3,000 K, followed by 5,000 cooling steps of 0.005 ps. 5,000 steps of energy minimization were applied to produce the final structures, which were recorded as coordinate files. The resulting structures had no distance or dihedral angle violation greater than 0.3 Å or 5°, respectively. The fifteen lowest energy structures were chosen for visualization and statistical analyses. Structure evaluation was performed with the program PROCHECK-NMR^102^; the percentage of residues in the most favored, additionally allowed, generously allowed and disallowed regions is 95.4, 4.5, 0.1 and 0.0, respectively. Visualization was performed with MOLMOL^103^ and PyMOL (PyMOL Molecular Graphics System, http://www.pymol.org). The electrostatic surface of E6AP AZUL was generated by the Poisson-Boltzmann (APBS) method^104, 105^.

## Acknowledgements

This work was supported by the Intramural Research Program through the CCR, NCI, NIH (1 ZIA BC011490).

## Author contributions

G.R.B. performed molecular biology and cell biology studies. X.C. performed CD, NMR and structure calculations. R.C. designed and generated plasmids for making *ΔRAZUL* cells. M.O. and T.A. performed SPR. C.J. and T. A. performed mass spectrometry analysis. D.L.E. prepared cell lysate for GST pull-down. V.S. made and isolated clones of *ΔRAZUL* cells. S.G.T. performed ITC. N.I.T. synthesized peptides. K.J.W designed experiments. G.R.B., X.C. and K.J.W wrote/edited the manuscript.

## Data and code availability

Atomic coordinates for RAZUL:AZUL have been deposited in the Protein Data Bank (PDB) with accession number 6U19. Chemical shift assignments have been deposited in the Biological Magnetic Resonance Data Bank (BMRB) with accession number 27875.

## Reference

1. Ehlinger, A. & Walters, K. J. Structural insights into proteasome activation by the 19S regulatory particle. Biochemistry 52, 3618–3628, doi:10.1021/bi400417a (2013).

2. Finley, D., Chen, X. & Walters, K. J. Gates, Channels, and Switches: Elements of the Proteasome Machine. Trends Biochem. Sci. 41, 77–93, doi:10.1016/j.tibs.2015.10.009 (2016).

3. Randles, L. & Walters, K. J. Ubiquitin and its binding domains. Front Biosci (Landmark Ed) 17, 2140–2157 (2012).

4. Deshaies, R. J. & Joazeiro, C. A. RING domain E3 ubiquitin ligases. Annual review of biochemistry 78, 399–434, doi:10.1146/annurev.biochem.78.101807.093809 (2009).

5. Rotin, D. & Kumar, S. Physiological functions of the HECT family of ubiquitin ligases. Nature reviews. Molecular cell biology 10, 398–409, doi:10.1038/nrm2690 (2009).

6. Scheffner, M. & Kumar, S. Mammalian HECT ubiquitin-protein ligases: biological and pathophysiological aspects. Biochimica et biophysica acta 1843, 61–74, doi:10.1016/j.bbamcr.2013.03.024 (2014).

7. Elsasser, S. et al. Proteasome subunit Rpn1 binds ubiquitin-like protein domains. Nat. Cell Biol. 4, 725--730, doi:10.1038/ncb845 (2002).

8. Hiyama, H. et al. Interaction of hHR23 with S5a. The ubiquitin-like domain of hHR23 mediates interaction with S5a subunit of 26 S proteasome. J. Biol. Chem. 274, 28019-28025 (1999).

9. Husnjak, K. et al. Proteasome subunit Rpn13 is a novel ubiquitin receptor. Nature 453, 481–488, doi:10.1038/nature06926 (2008).

10. Schreiner, P. et al. Ubiquitin docking at the proteasome through a novel pleckstrin-homology domain interaction. Nature 453, 548–552, doi:10.1038/nature06924 (2008).

11. Shi, Y. et al. Rpn1 provides adjacent receptor sites for substrate binding and deubiquitination by the proteasome. Science 351, doi:10.1126/science.aad9421 (2016).

12. Young, P., Deveraux, Q., Beal, R. E., Pickart, C. M. & Rechsteiner, M. Characterization of two polyubiquitin binding sites in the 26 S protease subunit 5a. The Journal of biological chemistry 273, 5461–5467 (1998).

13. Walters, K. J., Kleijnen, M. F., Goh, A. M., Wagner, G. & Howley, P. M. Structural studies of the interaction between ubiquitin family proteins and proteasome subunit S5a. Biochemistry 41, 1767–1777 (2002).

14. Chen, X. et al. Structure of hRpn10 Bound to UBQLN2 UBL Illustrates Basis for Complementarity between Shuttle Factors and Substrates at the Proteasome. Journal of molecular biology 431, 939–955, doi:10.1016/j.jmb.2019.01.021 (2019).

15. Bertolaet, B. L. et al. UBA domains of DNA damage-inducible proteins interact with ubiquitin. Nat. Struct. Biol. 8, 417–422, doi:10.1038/87575 [pii] (2001).

16. Wilkinson, C. R. et al. Proteins containing the UBA domain are able to bind to multi-ubiquitin chains. Nat. Cell. Biol. 3, 939–943, doi:10.1038/ncb1001-939 [pii] (2001).

17. Wang, Q., Goh, A. M., Howley, P. M. & Walters, K. J. Ubiquitin recognition by the DNA repair protein hHR23a. Biochemistry 42, 13529–13535, doi:10.1021/bi035391j (2003).

18. Verma, R. et al. Role of Rpn11 metalloprotease in deubiquitination and degradation by the 26S proteasome. Science 298, 611–615, doi:10.1126/science.1075898 [pii] (2002).

19. Lam, Y. A., Xu, W., DeMartino, G. N. & Cohen, R. E. Editing of ubiquitin conjugates by an isopeptidase in the 26S proteasome. Nature 385, 737–740, doi:10.1038/385737a0 (1997).

20. Verma, R. et al. Proteasomal proteomics: identification of nucleotide-sensitive proteasome-interacting proteins by mass spectrometric analysis of affinity-purified proteasomes. Mol Biol Cell 11, 3425–3439 (2000).

21. Borodovsky, A. et al. A novel active site-directed probe specific for deubiquitylating enzymes reveals proteasome association of USP14. The EMBO journal 20, 5187–5196, doi:10.1093/emboj/20.18.5187 (2001).

22. Leggett, D. S. et al. Multiple associated proteins regulate proteasome structure and function. Mol. Cell 10, 495--507, doi:10.1016/S1097-2765(02)00638-X (2002).

23. Smith, D. M. et al. Docking of the proteasomal ATPases’ carboxyl termini in the 20S proteasome’s alpha ring opens the gate for substrate entry. Molecular cell 27, 731–744, doi:S1097-2765(07)00445-5 [pii] 10.1016/j.molcel.2007.06.033 (2007).

24. Gillette, T. G., Kumar, B., Thompson, D., Slaughter, C. A. & DeMartino, G. N. Differential roles of the COOH termini of AAA subunits of PA700 (19 S regulator) in asymmetric assembly and activation of the 26 S proteasome. J. Biol. Chem. 283, 31813–31822, doi:M805935200 [pii] 10.1074/jbc.M805935200 (2008).

25. Rabl, J. et al. Mechanism of gate opening in the 20S proteasome by the proteasomal ATPases. Molecular cell 30, 360–368, doi:S1097-2765(08)00175-5 [pii] 10.1016/j.molcel.2008.03.004 (2008).

26. Kane, R. C., Bross, P. F., Farrell, A. T. & Pazdur, R. Velcade: U.S. FDA approval for the treatment of multiple myeloma progressing on prior therapy. Oncologist 8, 508–513 (2003).

27. Herndon, T. M., et al. U.s. Food and Drug Administration approval: carfilzomib for the treatment of multiple myeloma. Clin Cancer Res 19, 4559–4563, doi:10.1158/1078-0432.CCR-13-0755 (2013).

28. Shirley, M. Ixazomib: First Global Approval. Drugs 76, 405–411, doi:10.1007/s40265-016-0548-5 (2016).

29. Anchoori, R. K. et al. A bis-benzylidine piperidone targeting proteasome ubiquitin receptor RPN13/ADRM1 as a therapy for cancer. Cancer cell 24, 791–805, doi:10.1016/j.ccr.2013.11.001 (2013).

30. Trader, D. J., Simanski, S. & Kodadek, T. A Reversible and Highly Selective Inhibitor of the Proteasomal Ubiquitin Receptor Rpn13 Is Toxic To Multiple Myeloma Cells. J. Am. Chem. Soc., doi:10.1021/jacs.5b02069 (2015).

31. Randles, L., Anchoori, R. K., Roden, R. B. & Walters, K. J. Proteasome Ubiquitin Receptor hRpn13 and its Interacting Deubiquitinating Enzyme Uch37 are Required for Proper Cell Cycle Progression. J. Biol. Chem., doi:10.1074/jbc.M115.694588 (2016).

32. Soong, R. S. et al. RPN13/ADRM1 inhibitor reverses immunosuppression by myeloid-derived suppressor cells. Oncotarget 7, 68489–68502, doi:10.18632/oncotarget.12095 (2016).

33. Anchoori, R. K. et al. Covalent Rpn13-Binding Inhibitors for the Treatment of Ovarian Cancer. ACS Omega 3, 11917–11929, doi:10.1021/acsomega.8b01479 (2018).

34. Song, Y. et al. Targeting proteasome ubiquitin receptor Rpn13 in multiple myeloma. Leukemia 30, 1877–1886, doi:10.1038/leu.2016.97 (2016).

35. Huibregtse, J. M., Scheffner, M. & Howley, P. M. Cloning and expression of the cDNA for E6-AP, a protein that mediates the interaction of the human papillomavirus E6 oncoprotein with p53. Molecular and cellular biology 13, 775–784 (1993).

36. Huibregtse, J. M., Scheffner, M. & Howley, P. M. Localization of the E6-AP regions that direct human papillomavirus E6 binding, association with p53, and ubiquitination of associated proteins. Molecular and cellular biology 13, 4918–4927 (1993).

37. Scheffner, M., Huibregtse, J. M., Vierstra, R. D. & Howley, P. M. The HPV-16 E6 and E6-AP complex functions as a ubiquitin-protein ligase in the ubiquitination of p53. Cell 75, 495–505 (1993).

38. Kishino, T., Lalande, M. & Wagstaff, J. UBE3A/E6-AP mutations cause Angelman syndrome. Nat Genet 15, 70–73, doi:10.1038/ng0197-70 (1997).

39. Matsuura, T. et al. De novo truncating mutations in E6-AP ubiquitin-protein ligase gene (UBE3A) in Angelman syndrome. Nat Genet 15, 74–77, doi:10.1038/ng0197-74 (1997).

40. Cooper, E. M., Hudson, A. W., Amos, J., Wagstaff, J. & Howley, P. M. Biochemical analysis of Angelman syndrome-associated mutations in the E3 ubiquitin ligase E6-associated protein. The Journal of biological chemistry 279, 41208–41217, doi:10.1074/jbc.M401302200 (2004).

41. Samaco, R. C., Hogart, A. & LaSalle, J. M. Epigenetic overlap in autism-spectrum neurodevelopmental disorders: MECP2 deficiency causes reduced expression of UBE3A and GABRB3. Hum Mol Genet 14, 483–492, doi:10.1093/hmg/ddi045 (2005).

42. Miao, S. et al. The Angelman syndrome protein Ube3a is required for polarized dendrite morphogenesis in pyramidal neurons. J Neurosci 33, 327–333, doi:10.1523/JNEUROSCI.2509-12.2013 (2013).

43. Avagliano Trezza, R., et al. Loss of nuclear UBE3A causes electrophysiological and behavioral deficits in mice and is associated with Angelman syndrome. Nat Neurosci 22, 1235–1247, doi:10.1038/s41593-019-0425-0 (2019).

44. Lemak, A., Yee, A., Bezsonova, I., Dhe-Paganon, S. & Arrowsmith, C. H. Zn-binding AZUL domain of human ubiquitin protein ligase Ube3A. Journal of biomolecular NMR 51, 185–190, doi:10.1007/s10858-011-9552-y (2011).

45. Kuhnle, S. et al. Angelman syndrome-associated point mutations in the Zn(2+)-binding N-terminal (AZUL) domain of UBE3A ubiquitin ligase inhibit binding to the proteasome. The Journal of biological chemistry 293, 18387–18399, doi:10.1074/jbc.RA118.004653 (2018).

46. Yi, J. J. et al. The autism-linked UBE3A T485A mutant E3 ubiquitin ligase activates the Wnt/beta-catenin pathway by inhibiting the proteasome. The Journal of biological chemistry 292, 12503–12515, doi:10.1074/jbc.M117.788448 (2017).

47. Martinez-Noel, G. et al. Identification and proteomic analysis of distinct UBE3A/E6AP protein complexes. Molecular and cellular biology 32, 3095–3106, doi:10.1128/MCB.00201-12 (2012).

48. Chu, B. W. et al. The E3 ubiquitin ligase UBE3C enhances proteasome processivity by ubiquitinating partially proteolyzed substrates. The Journal of biological chemistry 288, 34575–34587, doi:10.1074/jbc.M113.499350 (2013).

49. Wang, Q. & Walters, K. J. Chemical shift assignments of the (poly)ubiquitin-binding region of the proteasome subunit S5a. Journal of biomolecular NMR 30, 231–232 (2004).

50. Wang, Q., Young, P. & Walters, K. J. Structure of S5a bound to monoubiquitin provides a model for polyubiquitin recognition. J. Mol. Biol. 348, 727–739, doi:10.1016/j.jmb.2005.03.007 (2005).

51. Zhang, N. et al. Structure of the s5a:k48-linked diubiquitin complex and its interactions with rpn13. Mol. Cell 35, 280–290, doi:10.1016/j.molcel.2009.06.010 (2009).

52. Lu, X., Liu, F., Durham, S. E., Tarasov, S. G. & Walters, K. J. A High Affinity hRpn2-Derived Peptide That Displaces Human Rpn13 from Proteasome in 293T Cells. PLoS One 10, e0140518, doi:10.1371/journal.pone.0140518 (2015).

53. Lu, X. et al. Structure of the Rpn13-Rpn2 complex provides insights for Rpn13 and Uch37 as anticancer targets. Nat Commun 8, 15540, doi:10.1038/ncomms15540 (2017).

54. Glickman, M. H. et al. A subcomplex of the proteasome regulatory particle required for ubiquitin-conjugate degradation and related to the COP9-signalosome and eIF3. Cell 94, 615–623, doi:10.1016/s0092-8674(00)81603-7 (1998).

55. Walters, K. J. et al. Characterizing protein-protein complexes and oligomers by nuclear magnetic resonance spectroscopy. Methods in enzymology 339, 238–258 (2001).

56. Chen, X. & Walters, K. J. Identifying and studying ubiquitin receptors by NMR. Methods Mol. Biol. 832, 279–303, doi:10.1007/978-1-61779-474-2_20 (2012).

57. Holm, L. Benchmarking Fold Detection by DaliLite v.5. Bioinformatics, doi:10.1093/bioinformatics/btz536 (2019).

58. Sims, J. J. & Cohen, R. E. Linkage-specific avidity defines the lysine 63-linked polyubiquitin-binding preference of rap80. Molecular cell 33, 775–783, doi:10.1016/j.molcel.2009.02.011 (2009).

59. Walters, K. J. & Chen, X. Measuring ubiquitin chain linkage: Rap80 uses a molecular ruler mechanism for ubiquitin linkage specificity. The EMBO journal 28, 2307–2308, doi:10.1038/emboj.2009.221 (2009).

60. Hornbeck, P. V., et al. PhosphoSitePlus, 2014: mutations, PTMs and recalibrations. Nucleic acids research 43, D512–520, doi:10.1093/nar/gku1267 (2015).

61. Deveraux, Q., Ustrell, V., Pickart, C. & Rechsteiner, M. A 26 S protease subunit that binds ubiquitin conjugates. J. Biol. Chem. 269, 7059--7061 (1994).

62. Wright, P. E. & Dyson, H. J. Intrinsically disordered proteins in cellular signalling and regulation. Nature reviews. Molecular cell biology 16, 18–29, doi:10.1038/nrm3920 (2015).

63. Dyson, H. J. & Wright, P. E. Coupling of folding and binding for unstructured proteins. Curr Opin Struct Biol 12, 54–60 (2002).

64. Lee, H. et al. Local structural elements in the mostly unstructured transcriptional activation domain of human p53. The Journal of biological chemistry 275, 29426–29432, doi:10.1074/jbc.M003107200 (2000).

65. Vise, P. D., Baral, B., Latos, A. J. & Daughdrill, G. W. NMR chemical shift and relaxation measurements provide evidence for the coupled folding and binding of the p53 transactivation domain. Nucleic acids research 33, 2061–2077, doi:10.1093/nar/gki336 (2005).

66. Wells, M. et al. Structure of tumor suppressor p53 and its intrinsically disordered N-terminal transactivation domain. Proceedings of the National Academy of Sciences of the United States of America 105, 5762–5767, doi:10.1073/pnas.0801353105 (2008).

67. Kussie, P. H. et al. Structure of the MDM2 oncoprotein bound to the p53 tumor suppressor transactivation domain. Science 274, 948–953, doi:10.1126/science.274.5289.948 (1996).

68. Borcherds, W. et al. Disorder and residual helicity alter p53-Mdm2 binding affinity and signaling in cells. Nat Chem Biol 10, 1000–1002, doi:10.1038/nchembio.1668 (2014).

69. Hamazaki, J. et al. A novel proteasome interacting protein recruits the deubiquitinating enzyme UCH37 to 26S proteasomes. EMBO J. 25, 4524–4536, doi:10.1038/sj.emboj.7601338 (2006).

70. Qiu, X. B. et al. hRpn13/ADRM1/GP110 is a novel proteasome subunit that binds the deubiquitinating enzyme, UCH37. EMBO J. 25, 5742-5753, doi:10.1038/sj.emboj.7601450 (2006).

71. Yao, T. et al. Proteasome recruitment and activation of the Uch37 deubiquitinating enzyme by Adrm1. Nat. Cell. Biol. 8, 994–1002, doi:10.1038/ncb1460 (2006).

72. Lander, G. C. et al. Complete subunit architecture of the proteasome regulatory particle. Nature 482, 186–191, doi:10.1038/nature10774 (2012).

73. Lasker, K. et al. Molecular architecture of the 26S proteasome holocomplex determined by an integrative approach. Proc. Natl. Acad. Sci. U. S. A. 109, 1380–1387, doi:10.1073/pnas.1120559109 (2012).

74. Besche, H. C. et al. Autoubiquitination of the 26S proteasome on Rpn13 regulates breakdown of ubiquitin conjugates. The EMBO journal 33, 1159–1176, doi:10.1002/embj.201386906 (2014).

75. Jacobson, A. D., MacFadden, A., Wu, Z., Peng, J. & Liu, C. W. Autoregulation of the 26S proteasome by in situ ubiquitination. Mol Biol Cell 25, 1824–1835, doi:10.1091/mbc.E13-10-0585 (2014).

76. Gottlieb, C. D., Thompson, A. C. S., Ordureau, A., Harper, J. W. & Kopito, R. R. Acute unfolding of a single protein immediately stimulates recruitment of ubiquitin protein ligase E3C (UBE3C) to 26S proteasomes. The Journal of biological chemistry, doi:10.1074/jbc.RA119.009654 (2019).

77. Zuin, A. et al. Rpn10 monoubiquitination orchestrates the association of the ubiquilin-type DSK2 receptor with the proteasome. Biochem J 472, 353–365, doi:10.1042/BJ20150609 (2015).

78. Keren-Kaplan, T. et al. Structure of ubiquitylated-Rpn10 provides insight into its autoregulation mechanism. Nat Commun 7, 12960, doi:10.1038/ncomms12960 (2016).

79. Isasa, M. et al. Monoubiquitination of RPN10 regulates substrate recruitment to the proteasome. Molecular cell 38, 733–745, doi:10.1016/j.molcel.2010.05.001 (2010).

80. Crosas, B. et al. Ubiquitin chains are remodeled at the proteasome by opposing ubiquitin ligase and deubiquitinating activities. Cell 127, 1401–1413, doi:10.1016/j.cell.2006.09.051 (2006).

81. You, J. & Pickart, C. M. A HECT domain E3 enzyme assembles novel polyubiquitin chains. The Journal of biological chemistry 276, 19871–19878, doi:10.1074/jbc.M100034200 (2001).

82. Lu, Y., Lee, B.-h., King, R. W., Finley, D. & Kirschner, M. W. Substrate degradation by the proteasome: a single-molecule kinetic analysis. Science 348, 1250834, doi:10.1126/science.1250834 (2015).

83. Meyer, H. J. & Rape, M. Enhanced protein degradation by branched ubiquitin chains. Cell 157, 910–921, doi:10.1016/j.cell.2014.03.037 (2014).

84. O’Leary, C. E., Lewis, E. L. & Oliver, P. M. Ubiquitylation as a Rheostat for TCR Signaling: From Targeted Approaches Toward Global Profiling. Front Immunol 6, 618, doi:10.3389/fimmu.2015.00618 (2015).

85. Oda, H., Kumar, S. & Howley, P. M. Regulation of the Src family tyrosine kinase Blk through E6AP-mediated ubiquitination. Proceedings of the National Academy of Sciences of the United States of America 96, 9557–9562, doi:10.1073/pnas.96.17.9557 (1999).

86. Collins, J. M., Porter, K. A., Singh, S. K. & Vanier, G. S. High-efficiency solid phase peptide synthesis (HE-SPPS). Org Lett 16, 940–943, doi:10.1021/ol4036825 (2014).

87. Chen, X. et al. Structures of Rpn1 T1:Rad23 and hRpn13:hPLIC2 Reveal Distinct Binding Mechanisms between Substrate Receptors and Shuttle Factors of the Proteasome. Structure 24, 1257–1270, doi:10.1016/j.str.2016.05.018 (2016).

88. van Stokkum, I. H., Spoelder, H. J., Bloemendal, M., van Grondelle, R. & Groen, F. C. Estimation of protein secondary structure and error analysis from circular dichroism spectra. Anal Biochem 191, 110–118, doi:10.1016/0003-2697(90)90396-q (1990).

89. Whitmore, L. & Wallace, B. A. DICHROWEB, an online server for protein secondary structure analyses from circular dichroism spectroscopic data. Nucleic acids research 32, W668–673, doi:10.1093/nar/gkh371 (2004).

90. Whitmore, L. & Wallace, B. A. Protein secondary structure analyses from circular dichroism spectroscopy: methods and reference databases. Biopolymers 89, 392–400, doi:10.1002/bip.20853 (2008).

91. Abdul-Gader, A., Miles, A. J. & Wallace, B. A. A reference dataset for the analyses of membrane protein secondary structures and transmembrane residues using circular dichroism spectroscopy. Bioinformatics 27, 1630–1636, doi:10.1093/bioinformatics/btr234 (2011).

92. Kao, W. H., Beaudenon, S. L., Talis, A. L., Huibregtse, J. M. & Howley, P. M. Human papillomavirus type 16 E6 induces self-ubiquitination of the E6AP ubiquitin-protein ligase. J Virol 74, 6408–6417, doi:10.1128/jvi.74.14.6408-6417.2000 (2000).

93. Chari, R., Yeo, N. C., Chavez, A. & Church, G. M. sgRNA Scorer 2.0: A Species-Independent Model To Predict CRISPR/Cas9 Activity. ACS Synth Biol 6, 902–904, doi:10.1021/acssynbio.6b00343 (2017).

94. Ran, F. A. et al. Genome engineering using the CRISPR-Cas9 system. Nature protocols 8, 2281–2308, doi:10.1038/nprot.2013.143 (2013).

95. Delaglio, F. et al. NMRPipe: a multidimensional spectral processing system based on UNIX pipes. J. Biomol. NMR 6, 277–293 (1995).

96. Bartels, C., Xia, T. H., Billeter, M., Guntert, P. & Wuthrich, K. The program XEASY for computer-supported NMR spectral analysis of biological macromolecules. J. Biomol. NMR 6, 1–10, doi:10.1007/BF00417486 (1995).

97. Wishart, D. S. & Sykes, B. D. The 13C chemical-shift index: a simple method for the identification of protein secondary structure using 13C chemical-shift data. Journal of biomolecular NMR 4, 171–180 (1994).

98. Shen, Y., Delaglio, F., Cornilescu, G. & Bax, A. TALOS+: a hybrid method for predicting protein backbone torsion angles from NMR chemical shifts. J. Biomol. NMR 44, 213–223, doi:10.1007/s10858-009-9333-z (2009).

99. Schleucher, J. et al. A general enhancement scheme in heteronuclear multidimensional NMR employing pulsed field gradients. Journal of biomolecular NMR 4, 301–306 (1994).

100. Kay, L. E., Keifer, P. & Saarinen, T. Pure absorption gradient enhanced heteronuclear single quantum correlation spectroscopy with improved sensitivity. J. Am. Chem. Soc. 1992, 10663–10665 (1992).

101. Schwieters, C. D., Kuszewski, J. J., Tjandra, N. & Clore, G. M. The Xplor-NIH NMR molecular structure determination package. J. Magn. Reson. 160, 65–73 (2003).

102. Laskowski, R. A., Rullmannn, J. A., MacArthur, M. W., Kaptein, R. & Thornton, J. M. AQUA and PROCHECK-NMR: programs for checking the quality of protein structures solved by NMR. J. Biomol. NMR 8, 477–486 (1996).

103. Koradi, R., Billeter, M. & Wuthrich, K. MOLMOL: a program for display and analysis of macromolecular structures. J Mol Graph 14, 51–55, 29-32 (1996).

104. Baker, N. A., Sept, D., Joseph, S., Holst, M. J. & McCammon, J. A. Electrostatics of nanosystems: application to microtubules and the ribosome. Proceedings of the National Academy of Sciences of the United States of America 98, 10037–10041, doi:10.1073/pnas.181342398 (2001).

105. Dolinsky, T. J., Nielsen, J. E., McCammon, J. A. & Baker, N. A. PDB2PQR: an automated pipeline for the setup of Poisson-Boltzmann electrostatics calculations. Nucleic acids research 32, W665–667, doi:10.1093/nar/gkh381 (2004).

