## Supplemental Files for "Structure of E3 ligase E6AP with a novel proteasome-binding site provided by substrate receptor hRpn10"

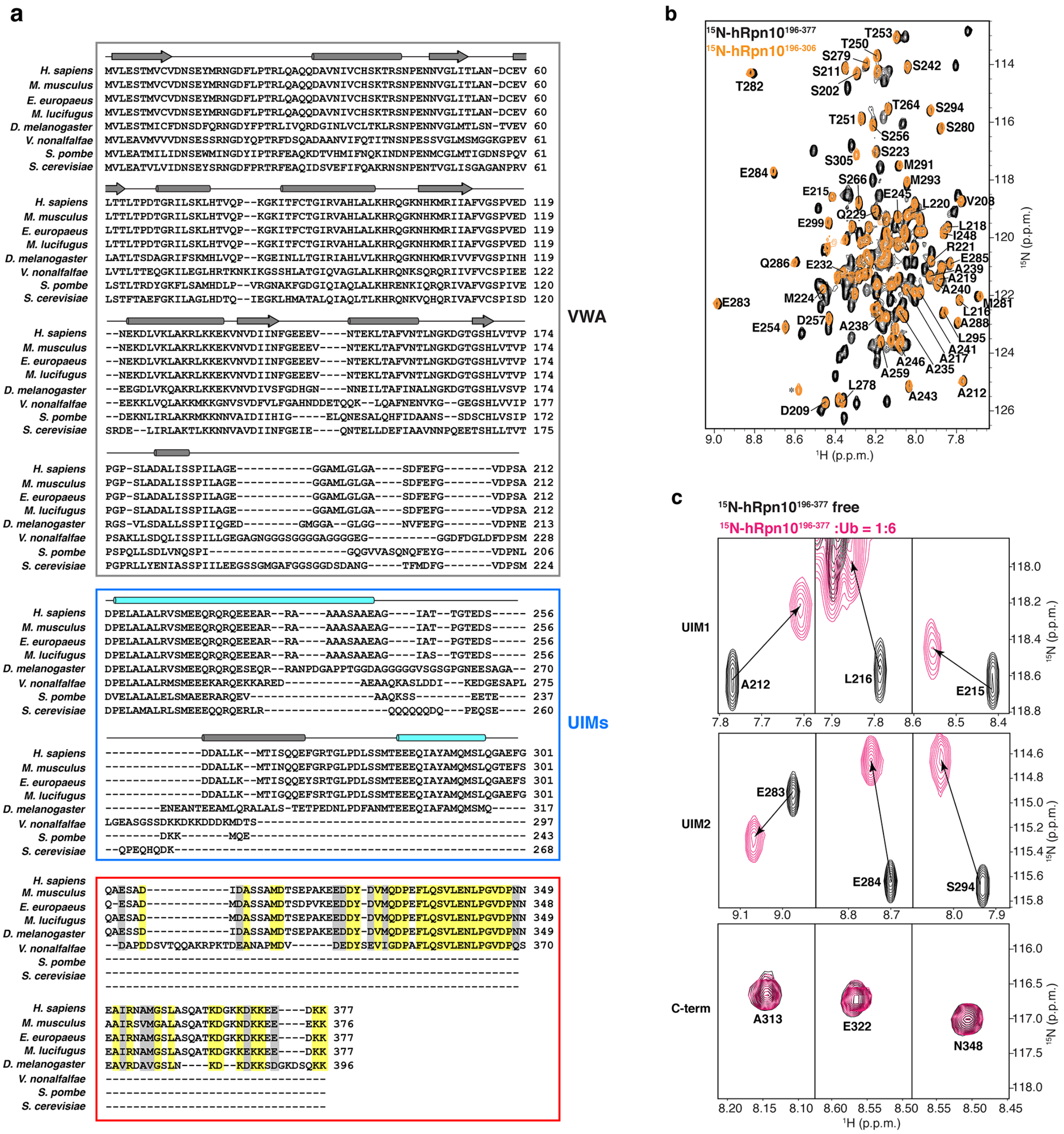


**Supplementary Figure 1. Rpn10 C-terminus is conserved in higher eukaryotes and does not interact with hRpn10 UIM region or ubiquitin.** (**a**) Sequence alignment of Rpn10 homologues from *H. sapiens* (P55036), *M. musculus* (O35226), *E. europaeus* (A0A1S3A7R2), *M. lucifugus* (G1PRQ7), *D. melanogaster* (P55035), *V. nonalfalfae* (A0A3M9Y9K5), *S. pombe* (O94444), *S. cerevisiae* (P38886) by *ClustalW*. Secondary structure elements of Rpn10 VWA (grey box) and the ubiquitin-binding UIM1- and UIM2-containing region (blue box) are indicated by arrows (β-strand) and cylinders (α-helix).

Identical and chemically similar residues in the Rpn10 C-terminal region (red box) are highlighted against a yellow and gray background respectively. (**b**) A selected region from $^{1}H$, $^{15}N$ HSQC spectra is superimposed for 0.2 mM $^{15}N$-hRpn10$^{196\text{-}377}$ (black) and $^{15}N$-hRpn10$^{196\text{-}306}$ (orange). Dispersed signals from UIM1 and UIM2 residues are labeled. A signal from the linker region between the N-terminal His-tag and hRpn10$^{196\text{-}306}$ sequence is indicated with an asterisk. **c** Selected regions from $^{1}H$, $^{15}N$ HSQC spectra of 0.2 mM $^{15}N$-hRpn10$^{196\text{-}377}$ (black) and with 6-fold molar excess unlabeled ubiquitin (pink). Amino acids from UIM1 (top, A212, E215, L216), UIM2 (middle, E283, E284, S294), and C-terminal region (bottom, A313, E322, N348) are included and labeled.

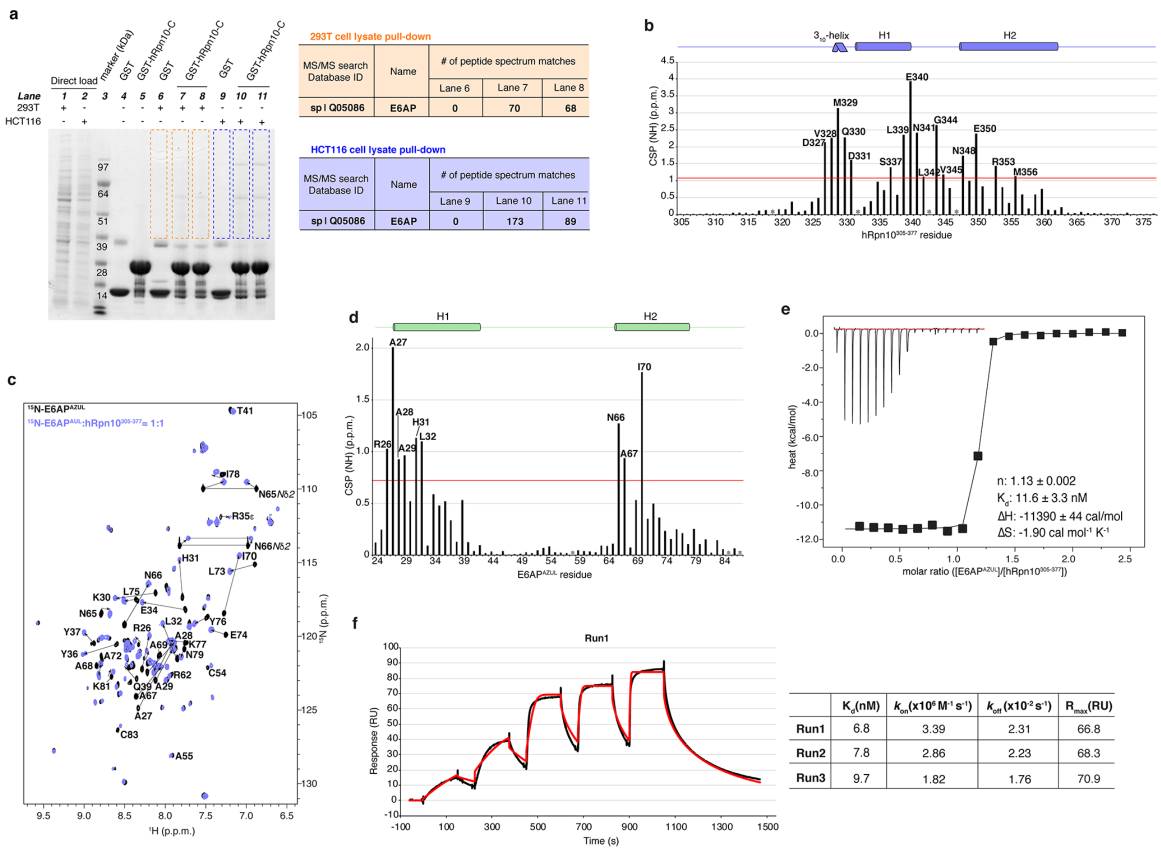


**Supplementary Figure 2. E6AP interacts with hRpn10$^{305\text{-}377}$ through its AZUL domain with high affinity.** (**a**) GST pull-down assay for GST-hRpn10$^{305\text{-}377}$ (GST-hRpn10-C) incubation with 293T (lane 7-8) or HCT116 (lane 10-11) cell lysate as indicated. GST protein was used as a control (lane 6 and 9). 293T or HCT116 cell lysate, GST-hRpn10$^{305\text{-}377}$, and GST protein were directly loaded in lane 1, 2, 4, and 5 as indicated. Each gel region of lane 6-11 above 39 kDa (boxed with orange or blue dashed lines) were cut into 12 bands, digested with trypsin, and analysed by mass spectrometry. The table shows the only identified hit for 293T or HCT116 lysate by searching MS/MS data against the Human Proteome database. (**b**) Chemical shift perturbation (CSP) plot for the data depicted in Figure 1C showing effects of 2-fold molar excess unlabeled E6AP AZUL addition to $^{15}$N-hRpn10$^{305\text{-}377}$. In **b** and **d**, the orange line indicates one standard deviation above the average value and prolines are indicated with grey asterisks. (**c**) $^{1}$H, $^{15}$N HSQC spectra of 0.2 mM $^{15}$N-E6AP AZUL (black) and with equimolar of unlabeled hRpn10$^{305\text{-}377}$ (blue).

E6AP AZUL signals that shift following addition of hRpn10$^{305\text{-}377}$ are labeled. (**d**) CSP plot for the data depicted in **c** showing effects of equimolar unlabeled hRpn10$^{305\text{-}377}$ addition to $^{15}$N-E6AP AZUL. (**e**) ITC analysis of hRpn10$^{305\text{-}377}$ and AZUL interaction. 0.462 mM E6AP$^{24\text{-}87}$ was injected into a calorimeter cell containing 0.0405 mM hRpn10$^{305\text{-}377}$ and the data were fit to a one-site binding mode with the indicated thermodynamic parameters. (**f**) SPR analysis of GST-hRpn10$^{305\text{-}377}$ and AZUL interaction. Experimental data (black) and the globally fit curve (red) for one SPR experiment is displayed (left) and a table included for kinetic and affinity analysis performed for three independent experiments (right).

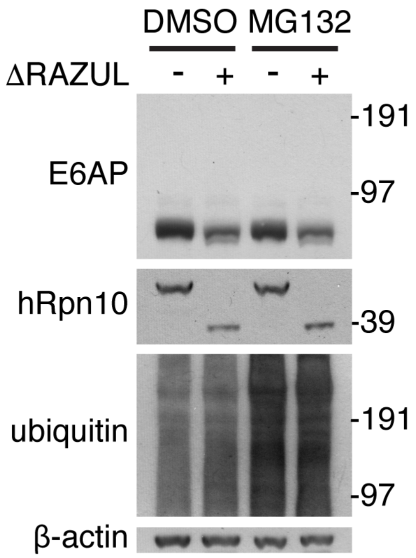


**Supplementary Figure 3. Reduced protein levels of E6AP and hRpn10 ΔRAZUL in *ΔRAZUL* cells are independent of proteasomal degradation.** Lysates from HCT116 (-) and ΔRAZUL clone14 (+) cells treated with 10 μM MG132 or DMSO (control) for 4 hours were immunoprobed for hRpn10, E6AP, ubiquitin, and β-actin (as a loading control).

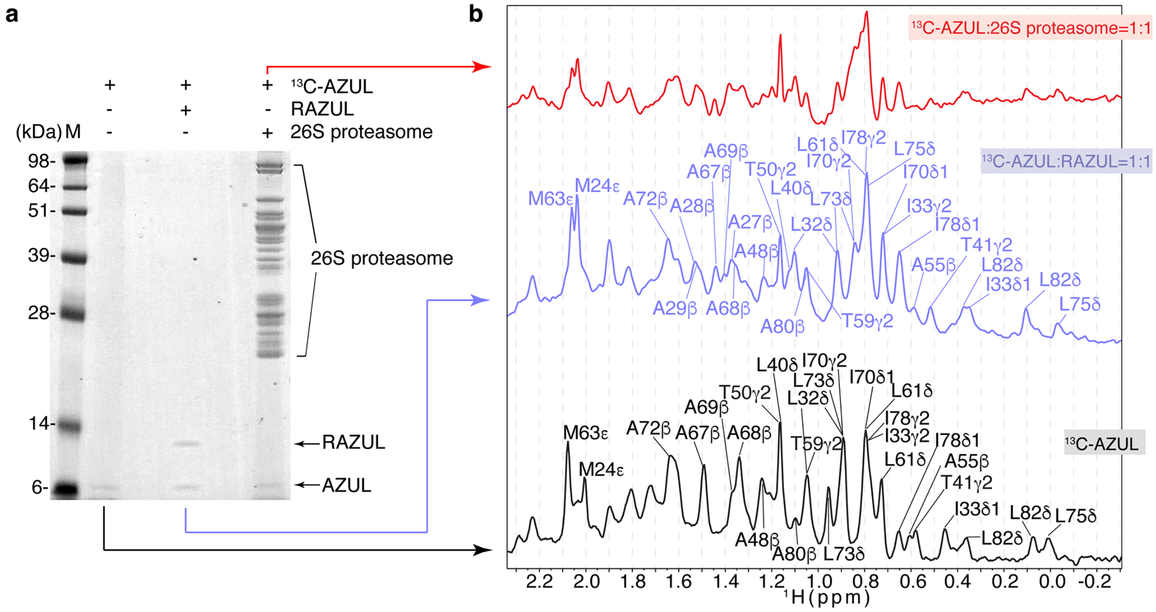


**Supplementary Figure 4. RAZUL:AZUL interaction was observed in the intact proteasome by using 1D $^{13}$C-edited, $^{1}$H NMR.** (**a**) SDS-PAGE gel for NMR samples of Fig. 3d including $^{13}$C-labeled AZUL alone and mixed with equimolar unlabeled RAZUL or with human 26S proteasome. (**b**) $^{13}$C-edited, $^{1}$H 1D NMR experiments acquired at 850 MHz, with a cryogenically cooled probe, and 25°C for free $^{13}$C-AZUL (black) or mixtures with equimolar unlabeled RAZUL (blue) or 26S proteasome (red). The concentration of each sample was 0.3 μM and 200,000 scans were recorded for each experiment.

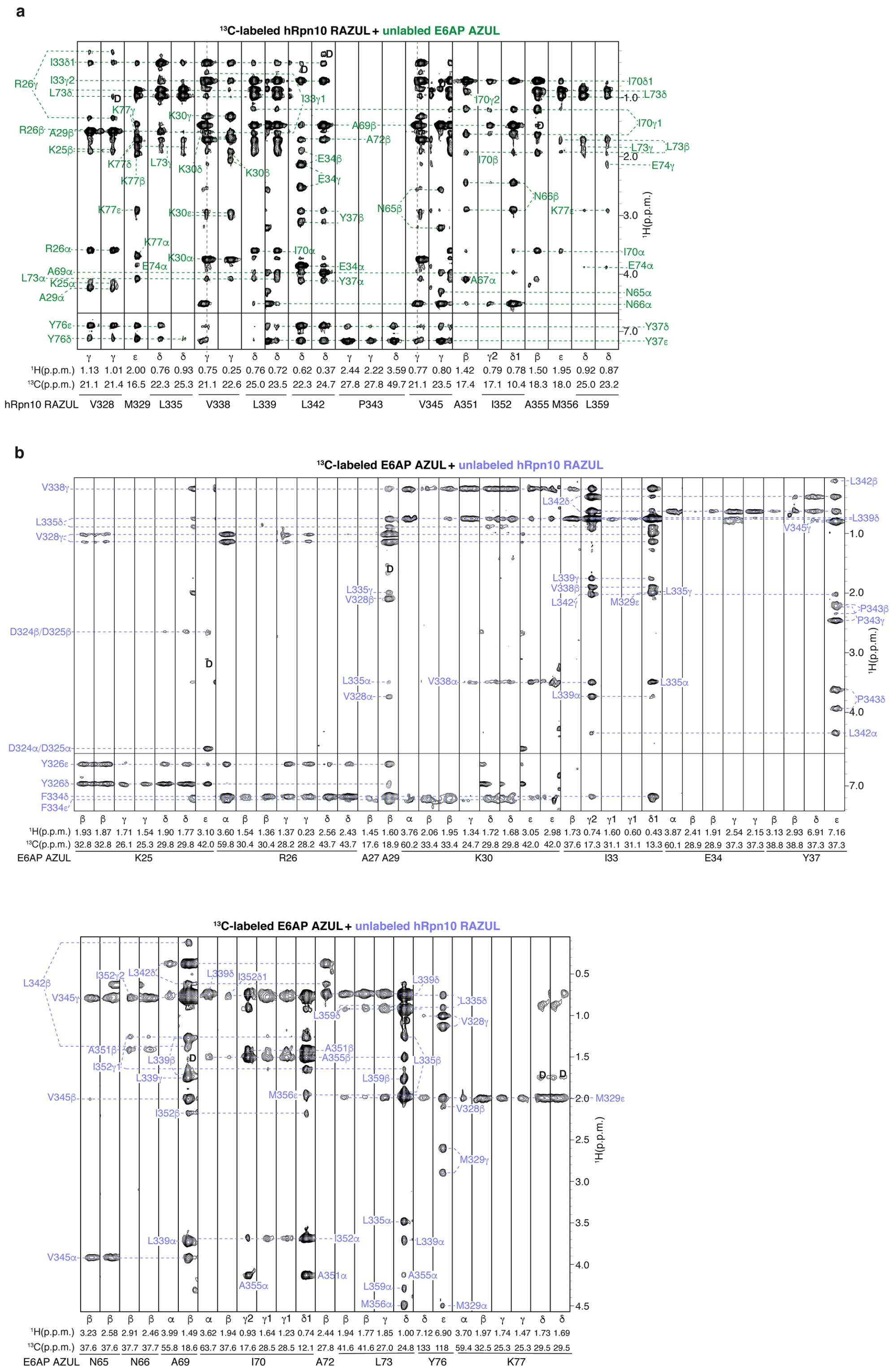

**Supplementary Figure 5. Intermolecular NOEs reflect the interaction of hRpn10 RAZUL and E6AP AZUL.** (**a**) Selected regions from a $^{13}$C-half-filtered NOESY experiment acquired with 0.5 mM $^{15}$N, $^{13}$C-labeled hRpn10 RAZUL mixed with unlabeled E6AP AZUL at 1.5-fold molar excess highlighting intermolecular NOE interactions. Labels inside (green) and outside (black) of the strips correspond to AZUL and RAZUL amino acids, respectively. (**b**) Selected regions from a $^{13}C$-half-filtered NOESY experiment acquired with 0.5 mM $^{15}N$, $^{13}C$-labeled E6AP AZUL mixed with unlabeled hRpn10 RAZUL at 1.5-fold molar excess highlighting intermolecular NOE interactions. Labels inside (blue) and outside (black) of the strips correspond to RAZUL and AZUL amino acids, respectively. Breakthrough diagonal peaks are labeled as ‘D’ in **a** and **b** and the experiments were each acquired with a 100 ms mixing time.

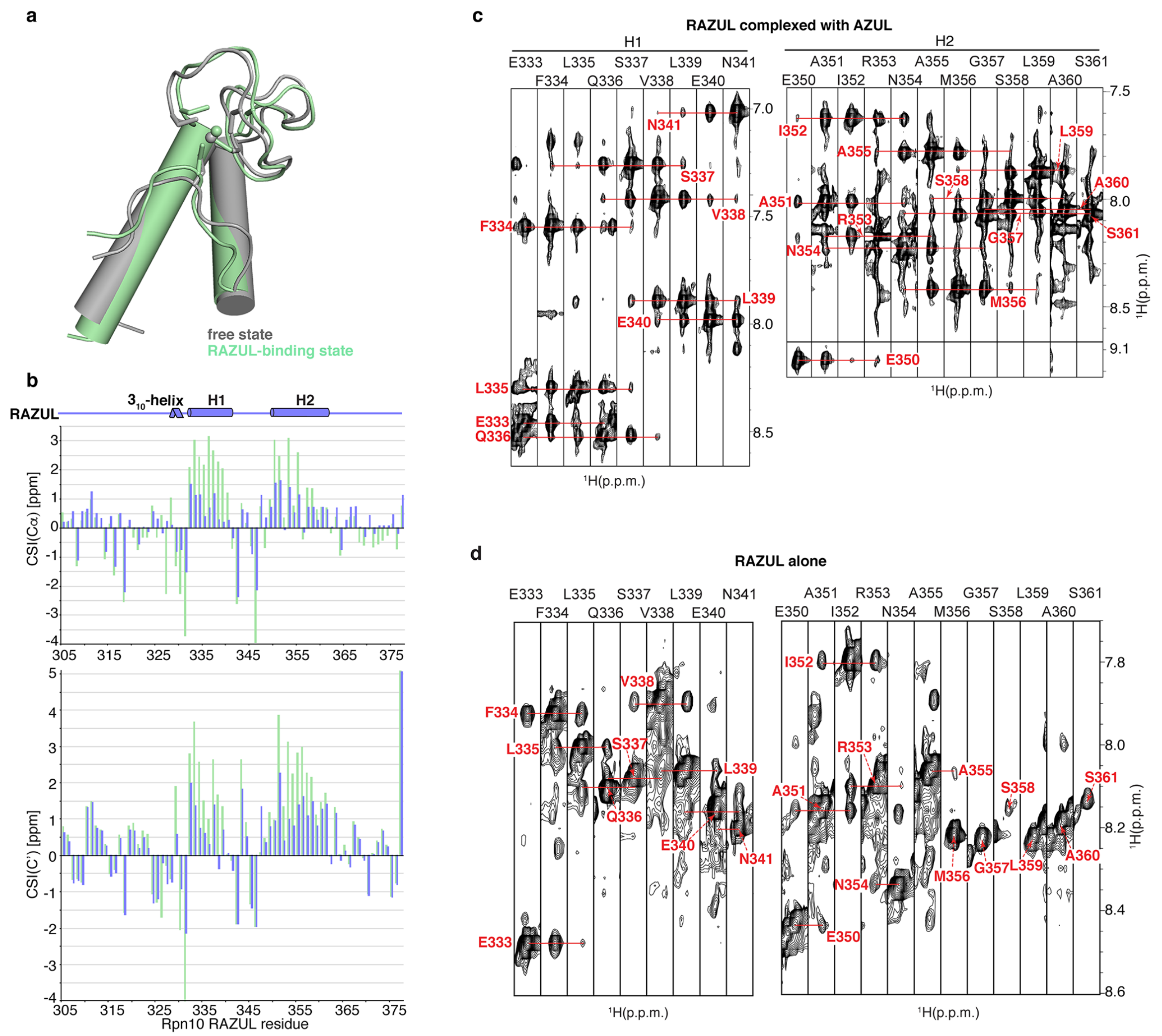


**Supplementary Figure 6. RAZUL switches from a poorly ordered state to a well-defined helical state upon binding AZUL.** (**a**) Superimposed structures of E6AP AZUL domain solved in its free state (grey, PDB 2KR1) and bound to RAZUL (green). Zn is displayed as a sphere in both structures. (**b**) The chemical shift index (CSI) value in part per million (ppm) for Cα (top panel) and C' (bottom panel) atoms is displayed for RAZUL alone (blue) and bound to AZUL (green). The RAZUL secondary structure is displayed in a cartoon at the top. (**c**) Selected $^{15}$N planes from a 3D $^{15}$N-dispersed NOESY spectrum (120 ms mixing time) recorded on a sample of 0.5 mM $^{15}$N, $^{13}$C-labeled hRpn10 RAZUL mixed with 1.5-fold molar excess unlabeled AZUL. (**d**) Selected $^{15}$N planes from a 3D $^{15}$N-dispersed NOESY spectrum recorded identically to **c** on a sample of 0.5 mM $^{15}$N, $^{13}$C-labeled hRpn10$^{196\text{-}377}$. In **c** and **d**, NOEs between backbone amide protons are highlighted by red connecting lines. All spectra were acquired on an 850 MHz spectrometer equipped with a cryogenically cooled probe and at 25°C.

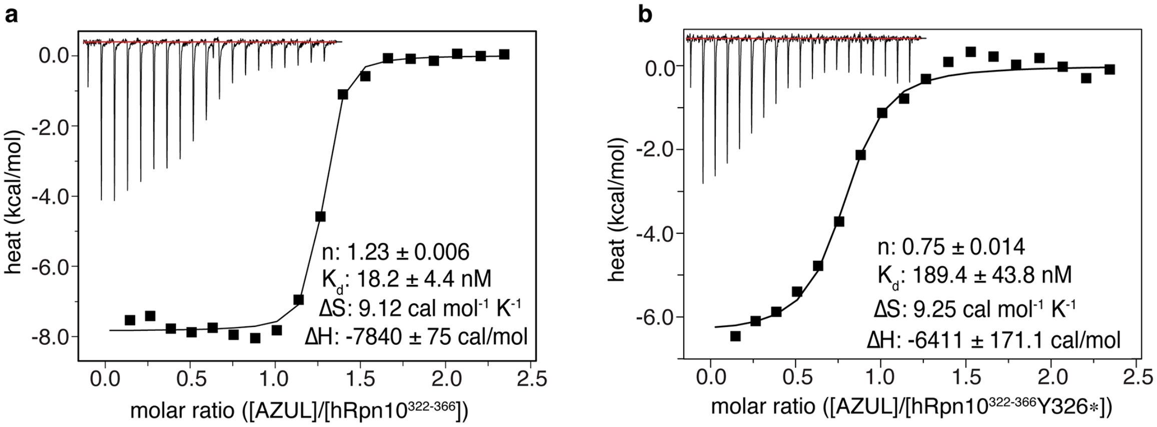


**Supplementary Figure 7. ITC analysis of AZUL interaction with hRpn10$^{322\text{-}366}$ without (a) and with Y326 phosphorylated (b)**. 0.110 mM E6AP$^{24\text{-}87}$ was injected into a calorimeter cell containing 0.01 mM hRpn10$^{322\text{-}366}$ without (**a**) or with Y326 phosphorylated (**b**) and the data were fit to a one-site binding mode with the indicated thermodynamic parameters.

**Supplementary Table 1.** sgRNA target site and oligonucleotides

| | | |
|---|---|---|
| Target Site | AGCAGGGCATCGTCTGAGTCTGG | TGCTAAGGTAGCAATGCTAATGG |
| Forward Oligo | CACCGAGCAGGGCATCGTCTGAGTC | CACCGTGCTAAGGTAGCAATGCTAA |
| Reverse Oligo | AAACGACTCAGACGATGCCCTGCTC | TTAGCATTGCTACCTTAGCAC |
